# Functional Dichotomy of Orbitofrontal Cortex and Anterior Cingulate Cortex Subregions in Decision-Making and Brain-Body Regulation

**DOI:** 10.1101/2024.11.26.625512

**Authors:** G.K. Papageorgiou, D.J. Gibson, K. Amemori, H.N. Schwerdt, M. Naim, M.C. Wang, T. Yoshida, J. Sharma, U. Upadhyay, G.R. Yang, A.M. Graybiel

## Abstract

Mood disorders are associated with complex disruptions in brain networks, including those associated with the orbitofrontal cortex (OFC) and pregenual anterior cingulate cortex (pACC). The functional contribution of the caudal OFC (cOFC) has remained largely unexplored. We investigated the functions of the cOFC and pACC in macaques performing an approach-avoidance task by the combination of multimodal recordings and electrical microstimulation (EMS) of the cOFC. We assessed neural, autonomic and behavioral responses. We found the cOFC to be sensitive to both positive and negative stimuli, whereas the pACC was singificantly more active during aversive outcomes. EMS of the cOFC increased avoidance behavior, suggesting a causal role for the OFC subdivision in cost-benefit decision-making. Physiological measurements were positively correlated with behavioral patterns, emphasizing body-brain synchronization during emotionally significant decision-making. We suggest that the cOFC contributes to inducing pessimistic states, thereby making its dysfunction a potential contributor to the etiology of mood disorders.

## Introduction

Mood-related disorders affect millions of individuals worldwide, leading to profound challenges in their emotional regulation and adaptive well-being. Central to many of these disorders is a pervasive negative affect and an inability to effectively manage stress and emotional balance. Dysfunctions within specific networks of cortical and noncortical regions are thought to underlie these symptoms (Chrysikou et al., 2022; Price & Drevets, 2010; Xu et al., 2021). Key among these are regions of the medial and ventral prefrontal neocortex, including the orbitofrontal cortex (OFC) and the pregenual anterior cingulate cortex (pACC), which in the normal state are thought to regulate cost-benefit decision-making, effortful behavior, and learning (Amemori et al., 2012; 2018; 2020; 2024; Price & Drevets, 2010; Rushworth & Behrens, 2008; Rushworth et al., 2011). The OFC is known for its extensive connections with sensory and its crucial role in evaluating rewards and punishments (Kringelbach & Rolls, 2004). Additionally, it functions in guiding adaptive behavior, allowing individuals to adjust their actions based on changing circumstances (Aupperle et al., 2015; Chau et al., 2015; Padoa-Schioppa & Assad, 2006; Papageorgiou et al., 2017). Similarly, the pACC has been implicated in a range of affective behaviors, with notable relevance to anxiety disorders and depression (Amemori et al., 2012; 2024; Ironside et al., 2020). These cortical regions are interconnected with the ventral and dorsal striatum and the amygdala and influence both emotional and motivational aspects of decision-making.

Whether the caudal part of the OFC (cOFC) contributes to these functions has been incompletely explored, despite its potential significance; it is situated at the limbic border of the neocortex (Aggleton et al., 2020; Barbas & Rempel-Clower, 1997; Murray & Fellows, 2022; Price & Drevets, 2010; Rempel-Clower & Barbas, 2000), and it has been shown to be an integral part of medial and prefrontal cortico-striato-limbic networks (Amemori et al., 2020; 2024; Eblen & Graybiel, 1995). This gap contrasts sharply with the well-mapped more rostral medial and lateral regions of the OFC (Ballesta et al., 2020; Fouragnan et al., 2019; Padoa-Schioppa & Assad, 2006; Papageorgiou et al., 2017). Potential clues to the functions of this extreme caudal zone of the OFC are provided by the similarities of its connections to those of the pACC (Amemori & Graybiel, 2012), notably that both receive relatively enhanced input from striosomes in the caudate nucleus and anterior putamen, themselves interconnected with regions of the limbic system. Moreover, both appear in concert in many human brain imaging studies (Volkow et al., 2012). Both the cOFC and other parts of the OFC have strong projections to the pACC, further implicating their role in mood regulation and decision-making processes.

We used these clues to guide a search for the response properties of cOFC neurons in macaque monkeys performing a modified version of the approach-avoidance (Ap-Av) task modified for non-human primates from one developed for healthy and clinical human populations (Amemori et al. 2012; Aupperle et al., 2015; Ironside et al., 2020) and in rodents for developing pharmaceutical drugs (Millan, 2003), and used in yet further modified form for understanding the decision-making processes (Friedman et al., 2015, 2017, 2020). Our goal here was to integrate the knowledge of the cOFC into the broader framework of cortico-cortical networks involved in decision-making and affective state. We hypothesized that the cOFC would have an instrumental function in underlying cost-benefit decision-making requiring acceptance or avoidance of combined good and bad offers and also in value-based learning, and that it could have causal effects on the animals’ observed behaviors. We combined multi-site recording in the cOFC and pACC and electrical microstimulation (EMS) of the cOFC with simultaneous collection of a series of physiological measurements.

The cOFC units exhibited balanced excitatory and inhibitory behavior during the cue period in which the cost-benefit offer was indicated, in contrast to pACC units simultaneously recorded, which tended to be mostly inhibitory, and further contrasts were evident. The EMS of the cOFC with either high (150-200μΑ) or low (5-15μΑ) currents induced increased avoidance behavior in the monkeys, indicating the causal role of cOFC in pessimistic bias in cost-benefit evaluation. Notably, we found significant correlations between the behavioral patterns of the monkeys and the autonomic and reaction time parameters measured during the decision-making task. This synchronization could hold cues to brain-body interactions contributing to mood disorders.

## Results

Two rhesus macaques, designated as subjects P and D, were trained over the course of approximately two years, during which they made the transition from a task-naive state to a level of task-expertise state in their use of directional saccades to one of two targets to report acceptance or rejection of a series of visually presented symbols representing compound offers of reward and punishment as in Amemori et al. (2012). The combined offers were delivered on a computer screen in front of the monkey and were composed of abutting horizontal bars whose 200x-independently variable lengths represented how much positive and negative reinforcement they would receive if they accepted the offer, where the rewarding juice was an incentive and aversive airpuff applied to the face was a deterrent. After the initial training regimen, we introduced the subjects to a conventional Ap-Av task illustrated in Fig. 1 (upper red-dotted rectangle).

**Figure 1.**
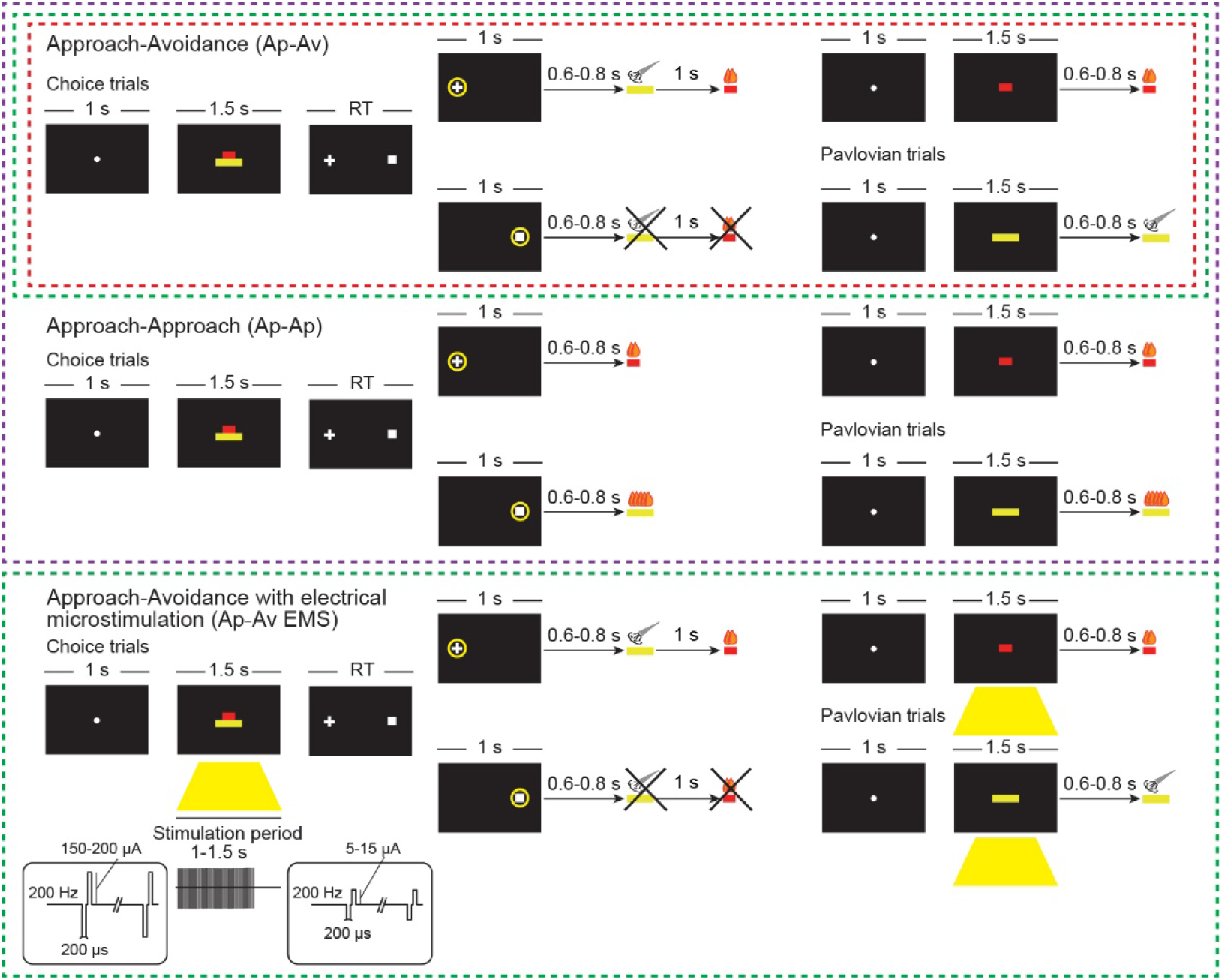
Experimental tasks. During the cue period of the Ap-Av task (red dotted rectangle), red and yellow horizontal bars signalled, respectively, the offered amounts of reward and punishment on the monitor. The monkeys made a decision between acceptance and rejection of the combined offer and reported this by choosing either of two targets (cross for acceptance/approach; square for rejection/avoidance) that appeared during the response period. The locations of the peripheral targets were alternated randomly. During the Pavlovian trials within the Ap-Av task, single cues were presented right after the fixation period, and the monkey experienced the respective positive or negative outcomes. During Ap-Av-Ap-Ap sessions (purple dotted rectangle), the Ap-Av task (upper diagram) was presented in the first and third blocks, and the Ap-Ap task (lower diagram) was presented in the second and fourth blocks. During the Ap-Ap cue period, the red and yellow horizontal bars on the monitor signaled two different offered amounts of reward. The monkeys made a decision between the two alternative options and reported this by choosing either of two targets (cross for choosing the red bar; square for choosing the yellow bar) that appeared during the response period. The locations of the targets were once again alternated randomly. During the Ap-Ap Pavlovian trials, single cues were presented right after the fixation period, and the monkey experienced the respective positive outcomes indicated by the length of either cue. During the Ap-Av EMS task (green dotted rectangles), the Ap-Av task was presented in the first and third blocks, and the Ap-Av EMS task was presented in the second block. During the Ap-Av EMS block, EMS was delivered simultaneously with the cue onset for 1 or 1.5 s of the cue period. The red and yellow horizontal bars, respectively signaling the offered amounts of reward and punishment, appeared on the monitor. The monkeys made a decision between acceptance and rejection of the combined offer and reported this by choosing either of two targets (cross for acceptance/approach; square for rejection/avoidance) that appeared during the response period. The locations of the targets were alternated randomly. During the Pavlovian trials, EMS was delivered as for the choice trials.

During daily (5 d/wk) sessions, the combination of two bars was presented to monkeys as the visual cue for decision-making. The cues consisted of a diverse array of approximately 40,000 combinations of reward and punishment, furnished randomly, to mimic the complexity inherent in real-world decision-making processes. In addition to the choice trials, we inserted Pavlovian trials, in which the animal experienced either positive or negative outcomes after single cues were presented. Pavlovian trials were introduced in 15% of the trials, interleaved in every 7 or 8 trials. In most cases, if a given Pavlovian trial were a punishment trial, then the next one would be a reward trial, and vice versa. In Pavlovian trials, after the fixation period, a single cue was presented on the screen, and the monkey needed only to fixate on the cue for the first 300 ms for the cue to stay on for the full 1.5-s presentation, followed by reinforcement or punishment. The fixation was considered successful as long as the eye position stayed within the 3° cue radius for the required time. Fixation breaks were allowed after 300 ms because otherwise the monkey would often exhibit fixation breaks to forfeit Pavlovian punishment trials. Due to the much smaller number of Pavlovian trials, and to investigate the parametric modulation of electrophysiological and physiological activity, we only presented bars of certain evenly spaced, preset sizes to ensure good coverage of the 200-interval distribution of rewards and punishments presented during the choice trials (see Methods). After this, the outcome followed a randomized delay uniformly distributed between 600-800 ms.

As a control decision-making task, we introduced an approach-approach (Ap-Ap) task that was largely identical to the Ap-Av task with the difference that both the red and yellow bars signaled reward amounts (Fig. 1; bottom half of purple dotted rectangle). The rewards offered by the red bar were given when the monkey chose to saccade at the cross, whereas the rewards offered by the yellow bar were given when the monkey chose to saccade on the square. As with the Ap-Av task, Pavlovian trials were presented, but now corresponding to the new contingencies. This experiment, was conducted in combination with the Ap-Av task, in each single session comprising two equal size blocks of the Ap-Av task interleaved with two equal size blocks of the Ap-Ap task, comprising the Ap-Av-Ap-Ap task (i.e., Ap-Av, Ap-Ap, Ap-Av, Ap-Ap). Finally, the animals were given the Ap-Av EMS task, which was identical to the Ap-Av task with the only difference being that we separated the task into three blocks of an equal number of trials (Fig. 1; green-dotted rectangle). In the second block, we microstimulated during the full cue period (1-1.5 s EMS) with either high (150-200 μΑ) or low (5-15 μA) currents, to determine whether and how varying levels of stimulation influence decision-making related to reward and punishment (Amemori et al., 2012; 2018; 2020; Ballesta et al., 2020). We note that previous research with the Ap-Av and Ap-Ap tasks were performed only under high currents (Amemori et al., 2012; 2018; 2020) and required monkeys to use their bodily somatic motor system to make decisions by reaching and pointing with a joystick to move a cursor on the screen. In the version of the experiments developed here, the monkeys were using saccades and did not have to make a change in reference frames to indicate their responses (Amemori et al., 2012; 2018; 2020). Here, the monkeys also encountered Pavlovian trials. We estimated the animals’ internal states during different task periods by recording lick rate, pupil diameter, reaction time and heart rate variability.

The animals executed 1050-1500 trials per recording session, and sessions lasted 4-6 hr. After the monkeys achieved mastery of the Ap-Av task, behavioral patterns like those depicted in Fig. 2a and b (top panel) emerged. For analysis, we included only sessions in which the decision boundaries had a positive slope (203 or 96.2% of all the sessions), indicating that the animals understood/performed well in the task, choosing an ‘optimal’ way to solve the task by accepting offers that more often came with larger rewards and smaller punishments (Fig. 2c). We excluded a small number of sessions (8 sessions or 3.8% of all the sessions) in which the decision boundaries had a negative slope. There was variability in the daily choices of the monkeys, partly because of the random combination of cues that were presented each day, and also due to the fact that on some occasions they rapidly decided to saccade to unfavorable side of the screen.

**Figure 2.**
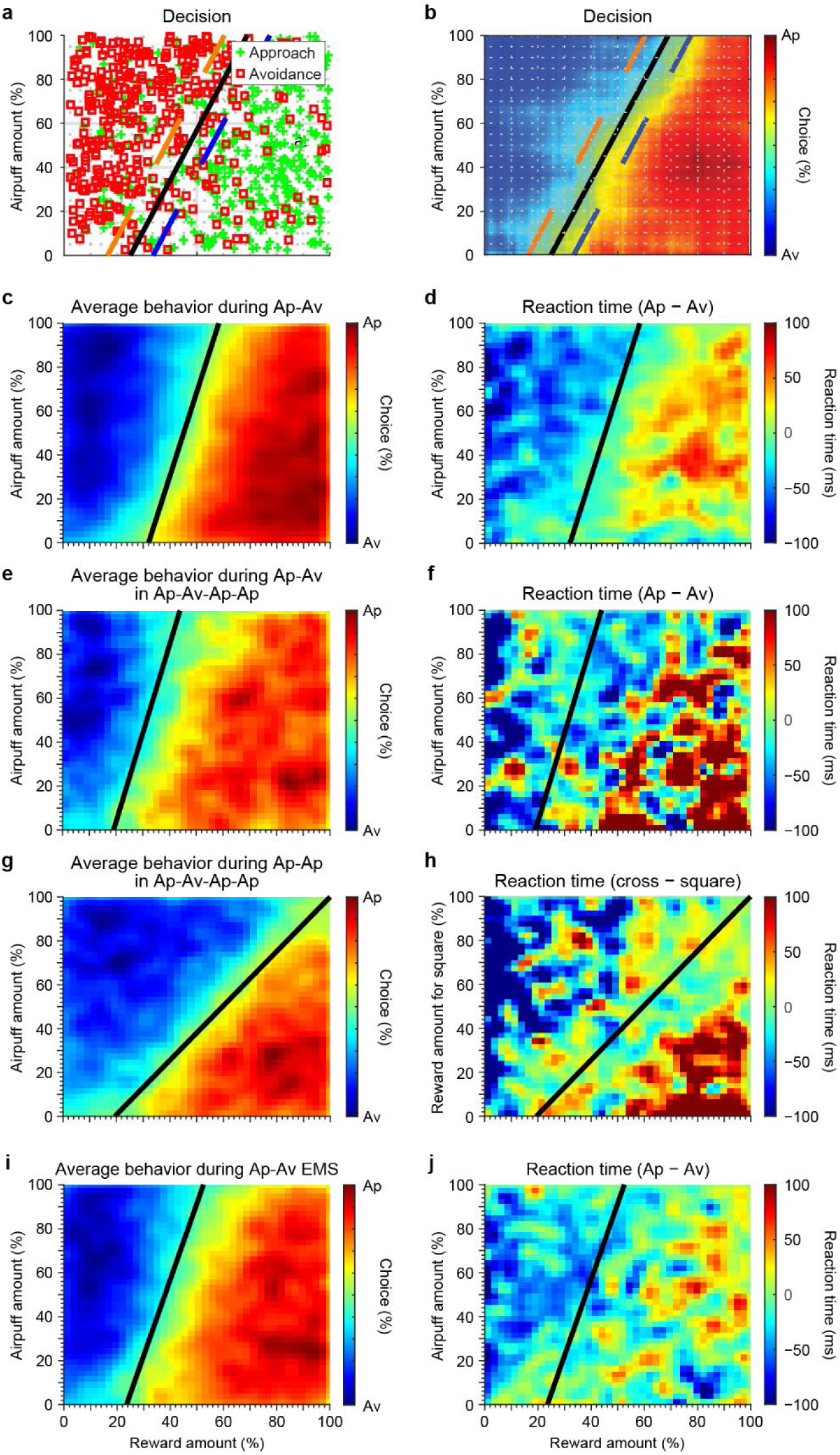
Physiological measurements across all experimental tasks. **a,b,** Session as an example of Ap-Av task performance with the original data (**a**) and a smoothed version of the same plot (**b**). The x-axis of these panels represents the percentage of reward (i.e., juice) offered, and the y-axis denotes the percentage of punishment (i.e., airpuff) offered. Trials in which subjects chose to approach (accept) the offers are marked with green crosses. Instances in which subjects opted to avoid (decline) the offers are denoted by red squares. The black lines indicate decision boundaries, which were mathematically derived through a linear regression model using Matlab’s ‘glmfit’ function (MathWorks, Natick, MA). The black line represents the 50% probability decision boundary, whereas the blue dashed line shows the 60% probability region, highlighting areas where approach behavior was more predominant. The orange dashed line marks the 40% probability region, underscoring areas where approach behavior was less predominant. **c,d,** Mean choice pattern (**c**) and difference in reaction times (**d**, see Methods) during Ap-Av task averaged across all sessions, showing a significant correlation between reaction times and observed choices (r = 0.697, p < 0.0001). **e,f,** Average choice pattern (**e**) and difference in reaction times (**f**) during the Ap-Av blocks in the Ap-Av-Ap-Ap task exhibit a positive and significant correlation (r = 0.18, p < 0.0001), highlighting optimal behavior. **g,h,** Average choice pattern (**g**) and difference in reaction times (**h**) during the Ap-Ap blocks in the Ap-Av-Ap-Ap task, with a significant positive correlation observed (r = 0.27, p < 0.0001). **i,j,** Average behavior (**i**) and difference in reaction times (**j**) in the Ap-Av EMS task, where behavior was again significantly correlated with reaction times (r = 0.47, p < 0.0001). The same decision boundaries used in the behavior plots (left column) have been applied to the reaction time plots (right column) for illustrative purposes.

We observed slower reaction times when animals approach better options (high reward, low airpuff) and reject worse options (high airpuff, low reward), likely reflecting a more deliberate decision-making process when the stakes were higher. This behavior suggests that animals take more time to evaluate the benefits of highly rewarding options, carefully weighing the potential gains before making an approach. Similarly, their hesitation quickly to reject highly aversive options could indicate a reluctance completely to abandon the decision, possibly due to lingering considerations of small rewards or the avoidance of regret. These slower responses highlight a cautious strategy aimed at maximizing decision accuracy in both favorable and unfavorable conditions, with animals exhibiting more cognitive control when faced with extreme trade-offs between reward and punishment (Fig. S1). We also calculated the difference in average reaction times (RT) between approach and avoidance trials across multiple sessions (see Methods). The difference in reaction times during the given session was significantly and positively correlated with the observed choices of the animals (Fig. 2d). During the Ap-Av blocks of the Ap-Av-Ap-Ap task, the animals exhibited near-optimal behavior, in which approach choices were positively and significantly correlated with the difference in RT (Fig. 2e,f). During the Ap-Ap blocks of the same session type, the animals also exhibited near-optimal behavior, choosing more often the largest of the two bars (Fig. 2g). Once again, the differences in RT were positively correlated with the decisions of the animals (Fig. 2h). Finally, during the Ap-Av EMS task, and their behavior was positively correlated once again with their differences in RT, demonstrating goal-directed decision-making (Fig. 2i,j).

### Neural activity during the Ap-Av task

We recorded 1,712 well-isolated single units from the cOFC (1,278 units) and pACC (434 units) of monkeys P and D (Fig. 3) with chronically implanted platinum iridium probes as well as acute recordings with Plexon S-probes, across all Ap-Av tasks (Fig. S2). We plotted the percentage of cOFC and pACC neurons that exhibited statistically significant changes in spike counts around various task events (chi-square test, p < 0.05, testing the change in spike counts 1 s before and 1 s after the onset of each event). The cOFC units showed a trend to be more active across most task events. This trend was particularly evident for the fixation point onset and Pavlovian red bar reward delivery onset. For the fixation point onset, there was a significant increase in cOFC activity (chi-square = 7.4782, p = 0.0062), with 54.37% of cOFC neurons showing significant activity compared to 37.35% in the pACC. Similarly, during the Pavlovian red bar reward delivery onset, there was also a notable rise in cOFC activity (chi-square = 6.6672, p = 0.0098), where 35.46% of cOFC neurons were significantly active, in contrast to 23.20% in the pACC. However, for two events, we observed the opposite pattern, specifically, during airpuff-related events: approach airpuff onset (Ap air ON) and Pavlovian airpuff onset (Yellow air ON). Among the events analyzed, the approach airpuff onset event exhibited a statistically significant difference in event-related unit counts between the cOFC (lower) and pACC (higher) (chi-square = 11.3735, p = 0.0007). The raw count of significantly active neurons was higher in the cOFC (670 neurons) than in the pACC (248 neurons), but this difference corresponded to 52.8% of the recorded neurons in the cOFC and 57.54% in the pACC. Additionally, during the Pavlovian yellow airpuff onset, there was a significant increase in pACC activity (chi-square = 5.0039, p = 0.0253), with 35.62% of pACC neurons showing significant activity, identical to the proportion observed during fixation point onset. These findings suggest that the cOFC is more actively involved during events related to reward processing and the initiation of trials, possibly linked to attentional and reward-evaluation processes. By contrast, the pACC shows greater involvement during events associated with aversive outcomes, such as the delivery of airpuffs following certain choices or cues. This pattern indicates that the pACC may play a significant role in processing the consequences of aversive events, contributing to learning and adaptation based on negative outcomes rather than responding directly to aversive stimuli.

**Figure 3.**
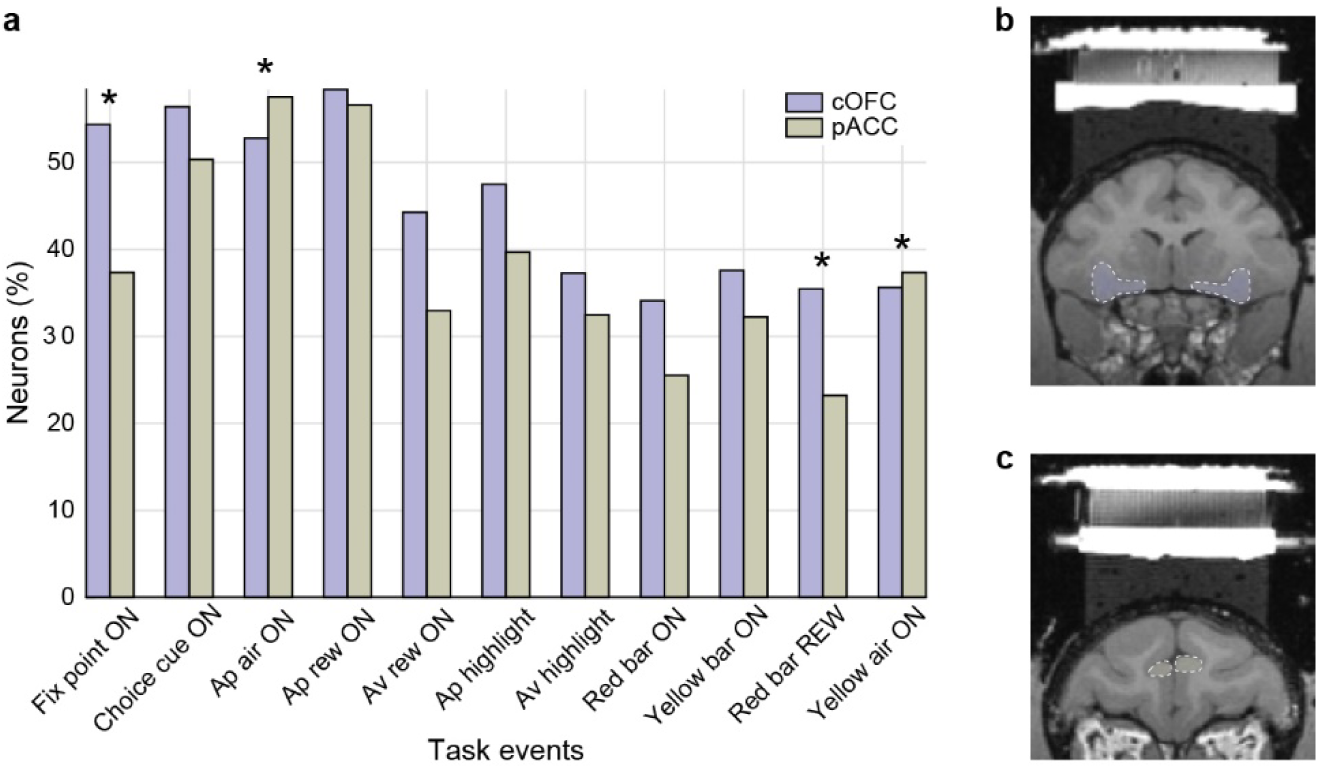
Event-related neural activity during Ap-Av. **a,** Neural activity by event and brain region during the Ap-Av task. The events plotted include the fixation point onset for all trials (Fix point ON), choice/offer cue onset (Choice cue ON), airpuff delivery onset during Ap choice trials (Ap air ON), reward delivery onset during Ap choice trials (Ap rew ON), small reward delivery onset during Av choice trials (Av rew ON), highlight around the cross for choosing Ap during choice trials (Ap highlight), highlight around the square for choosing Av during choice trials (Av highlight), Pavlovian trials reward cue onset (Red bar ON), Pavlovian trials airpuff cue onset (Yellow bar ON), Pavlovian trials reward delivery (Red bar REW), and Pavlovian trials airpuff delivery (Yellow air ON). *p < 0.05. **b,c,** Sample cOFC regions (**b**, shown in purple – white dotted outline) and sample pACC regions (**c**, shown in green – white dotted outline) from which the recordings have been performed.

We then used stepwise regression (Fig. 4), following the earlier protocol of Amemori et al.(2012), to determine whether the mean firing rates during the 1.5-s cue period were correlated with observable task variables: ‘Reward’ (the amount/duration of offered reward indicated by the length of the red bar), ‘Aversion’ (the amount/duration of offered airpuff indicated by the length of the yellow bar), the binary value of ‘Choice’ (Approach = 1, Avoidance = 0), the amount/duration of chosen reward as indicated in ‘Reward*Choice’, the amount/duration of chosen airpuff as indicated in ‘Airpuff*Choice’, and two hypothetical subjective variables, ‘Eutility’ (representing the expected utility of the offer) as well as ‘Conflict’ in decision-making. We also reasoned that if the cOFC or the pACC had a key role in integrating costs and benefits, this would likely be represented by both offered and chosen values.

**Figure 4.**
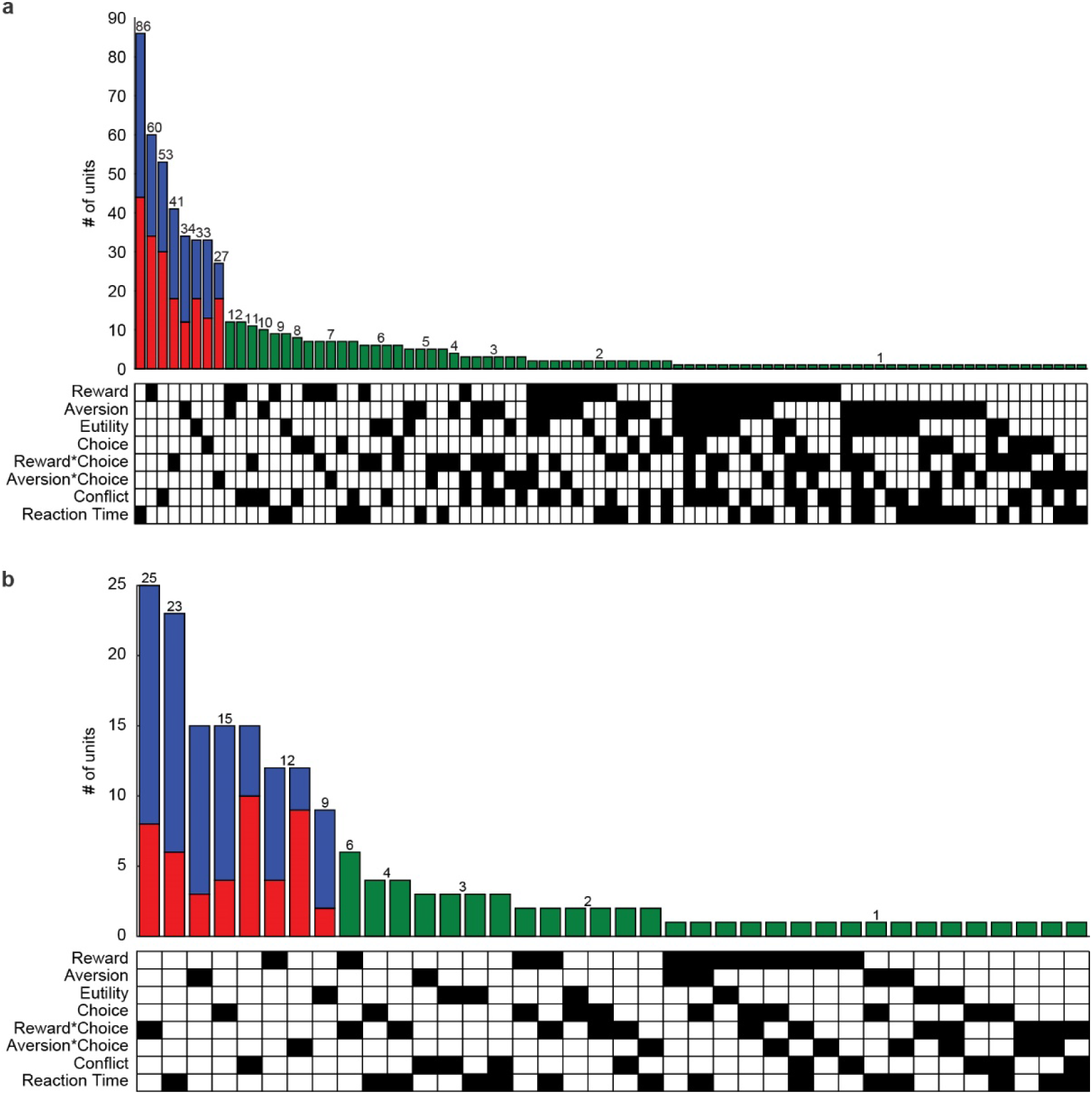
Classification of units recorded in the cOFC and pACC. Results of stepwise regression analysis for the Ap-Av task in the cOFC (**a**) and the pACC (**b**). Regression variables are reward size/duration (Reward), airpuff size/duration (Aversion), expected utility (Eutility), choice (Choice), chosen reward (Choice*Reward), chosen airpuff (Choice*Aversion), conflict in decisions (Conflict), and reaction times (RT). The histogram represents the number of units on the y-axis characterized by single (red-blue bars) or multiple (green bars) variables. These variables are then presented in the matrix below. The colored bars for the single variables demonstrate the positive (red) and negative (blue) correlations with the unit’s cue-related mean firing rates.

To further derive the subjective value of the chosen target, or the utility, we approximated the behavior using the conditional logit model (McFadden, 1974) and inferred utility inversely (Levy et al., 2010). We then used expected utility (Von Neumann & Morgenstern, 1947) as an explanatory variable corresponding to the continuous form of chosen value (Padoa-Schioppa, 2011). As in Amemori and Graybiel (2012), we used the entropy from the conditional logit model as an indication of decision conflict, a term we call ‘Conflict’ for the purposes of analysis. Stepwise regression (see Methods) indicated that activity during the decision period of the Ap-Av task was well characterized by linear combinations of the eight factors listed above (F-test, p < 0.05; Fig. 4). For the cOFC, the most significant predictor was Reaction Time (86 units), followed by Reward (60 units) and Conflict (53 units), with the remaining single predictors and several combinations of predictors following. By contrast, in the pACC, the most significant predictor was Reward*Choice (25 units), followed by Reaction Time (23 units), and then Aversion, Choice, and Conflict (15 units each). What stands out from this analysis is that the majority of the single predictors in the cOFC have a balanced positive/negative correlation with firing rates during the cue period, whereas in the pACC, these predictors are mostly negatively correlated with firing rates during the cue period.

### Differential task-related firing patterns in the pACC and cOFC

To facilitate comparison of the functional response properties of the pACC and the cOFC, we performed a heatmap analysis (Fig. 5). During the choice cue period (Fig. 5; Choice trials: top row), of the 428 pACC units analyzed, 76 (17.76%) were excitatory, 141 (32.95%) were inhibitory, and 211 (49.3%) were unresponsive. During the the airpuff delivery period in Approach choice trials, 85 (19.95%) units were excitatory, 163 (38.26%) were inhibitory, and 178 (41.78%) were unresponsive. During the reward delivery period in Approach choice trials, 178 (42.18%) units were excitatory, 57 (13.5%) were inhibitory, and 187 (44.3%) were unresponsive. In the small reward outcome period during the Av choice trials, 75 (17.73%) units were excitatory, 61 (14.42%) were inhibitory, and 287 (67.85%) were unresponsive.

**Figure 5.**
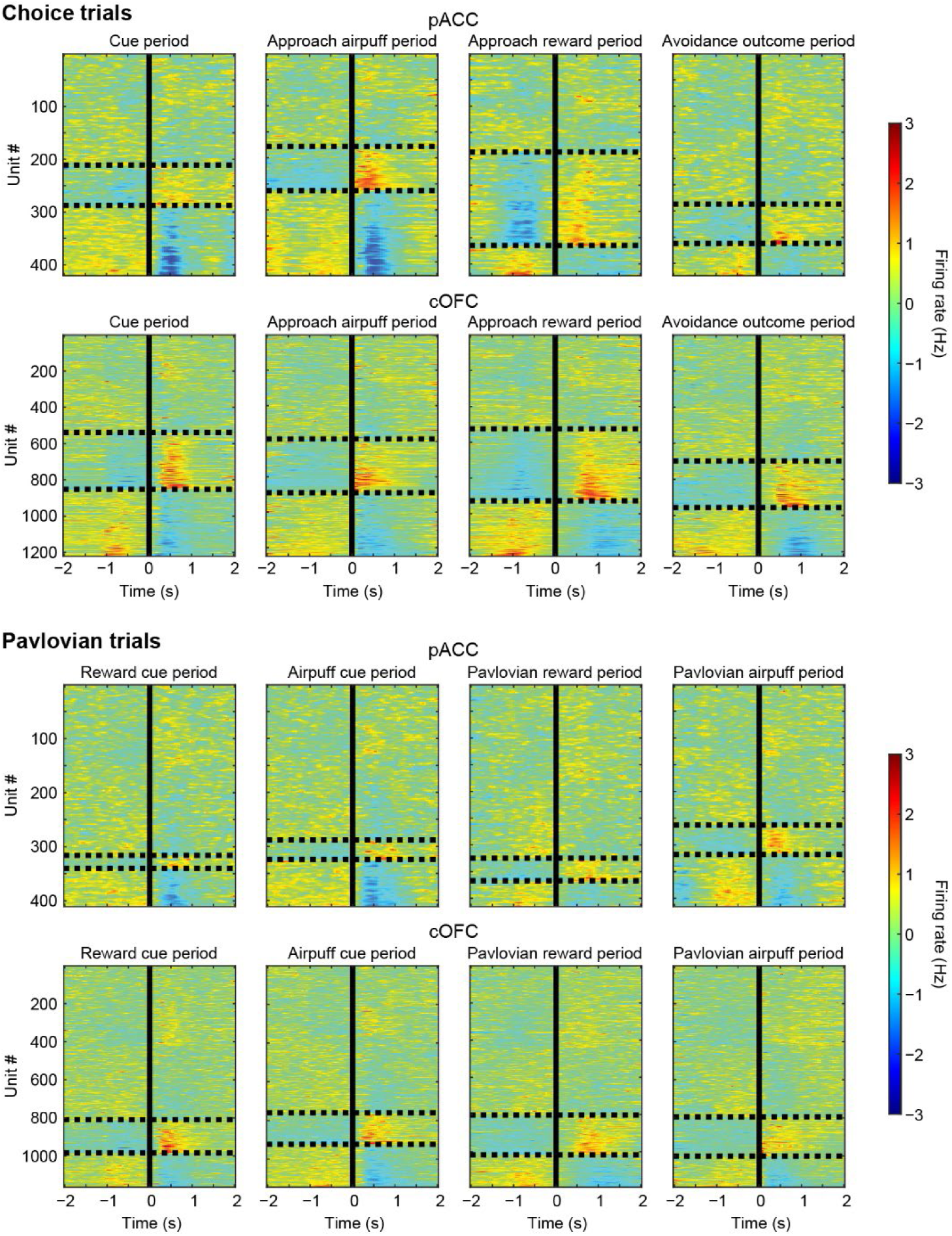
Differential firing patterns of pACC and cOFC units during choice and Pavlovian trials in the Ap-Av task. Each panel displays a color raster plot of the firing rates of individual units aligned on event onset, with three main compartments: the top compartment consists of units that were unresponsive to each event, the middle compartment (between the dotted lines) contains the excitatory units, and the bottom compartment contains the inhibitory units. The middle vertical line indicates the onset of each event. A 2-s period before and after the event onset is plotted. The y-axis of each panel represents the serial number of each unit. The top two rows show the results from the choice trials: the first row for pACC and the second row for cOFC. The bottom two rows present the results from the Pavlovian trials: the third row for pACC and the fourth row for cOFC. Statistical significance was tested using a chi-square test (p < 0.05) during the 1-s periods before and after the events. The heatmaps illustrate the variation in firing rates (in Hz), with warmer colors indicating higher firing rates and cooler colors indicating lower firing rates.

By contrast, in the cOFC (Fig. 5; Choice trials: second row), of the 1,257 units analyzed, 313 (24.94%) were excitatory in the cue period, 403 (32.11%) were inhibitory, and 539 (42.95%) were unresponsive. During Ap trials, specifically during the airpuff period, 303 (24.1%) units were excitatory, 367 (29.2%) were inhibitory, and 587 (46.7%) were unresponsive. During the reward delivery period in Ap trials, 402 (32.5%) units were excitatory, 309 (25%) were inhibitory, and 525 (42.48%) were unresponsive. During the Av outcome period, 258 (21.1%) units were excitatory, 269 (22%) were inhibitory, and 730 (56.98%) were unresponsive.

We further plotted the firing rates across all significant units in the cOFC and pACC during key Ap-Av task events. Around the cue onset event, we observed higher firing rates for the cOFC units than for the pACC units, with noticeable excitation after cue onset in the cOFC and slight inhibition during the same period in the pACC (Fig. S3a). For most task events, a larger proportion of cOFC units than pACC units exhibited excitatory responses, whereas the pACC had a greater proportion of inhibitory responses. This pattern highlights functional differences in how these brain regions process task-related information and suggests that the cOFC could be more involved in facilitating responses to stimuli, whereas the pACC could function more in modulating inhibitory control (Fig. S3b).

There was a similar pattern of firing rates during the Pavlovian trials, across both brain regions. Specifically, in the pACC, during the red cue period, 24 (5.85%) units were excitatory, 70 (17.1%) were inhibitory, and 316 (77.1%) were unresponsive. During the yellow cue period, 36 (8.76%) units were excitatory, 87 (21.17%) were inhibitory, and 288 (70.1%) were unresponsive. During the Pavlovian reward delivery period, 42 (10.12%) units were excitatory, 48 (11.57%) were inhibitory, and 325 (78.3%) were unresponsive. During the Pavlovian airpuff period, 55 (13.25%) units were excitatory, 97 (23.37%) were inhibitory, and 263 (63.37%) were unresponsive (Fig. 5).

Similarly, in the cOFC, during the red cue period, 175 (15.05%) units were excitatory, 179 (15.39%) were inhibitory, and 809 (69.56%) were unresponsive. During the yellow cue period, 163 (14.25%) units were excitatory, 221 (19.32%) were inhibitory, and 760 (66.43%) were unresponsive. During the Pavlovian reward delivery period, 214 (17.98%) units were excitatory, 173 (14.54%) were inhibitory, and 803 (67.48%) were unresponsive. Finally, during the Pavlovian airpuff period, 206 (17.71%) units were excitatory, 163 (14.02%) were inhibitory, and 794 (68.27%) were unresponsive (chi-square test, p < 0.05, during the 1-s period before and after events).

To determine how units in the two cortical regions responded when only the red bar or the yellow bar was presented during Pavlovian trials, we analyzed the effects of different cue/outcome sizes. We had six different sizes/durations of reward/punishment (i.e., 30, 60, 90, 120, 150, 180), designed to cover the range of sizes/durations (5-200) of the red/yellow bars in choice trials. We focused on four primary events of interest: the Reward Cue, which relates to the red bar onset; the Airpuff Cue, associated with the yellow bar onset; the reward delivery onset, which follows the presentation of the red bar; and the airpuff delivery onset, associated with the yellow bar’s presentation. For each brain region and event of interest, neuronal firing rates were plotted across time to visually assess changes in activity related to specific events (Fig. S4). For each event and brain region, we extracted neuronal activity data and calculated the mean firing rates within the specified time windows (i.e., 1 s before and after the onset of each event; see Fig. 6). To assess the significance of the observed trends, we performed linear regression on the bar values and calculated the t-statistic for the slope of the fitted line. The p-values for the slopes were computed to determine the significance of the trends.

**Figure 6.**
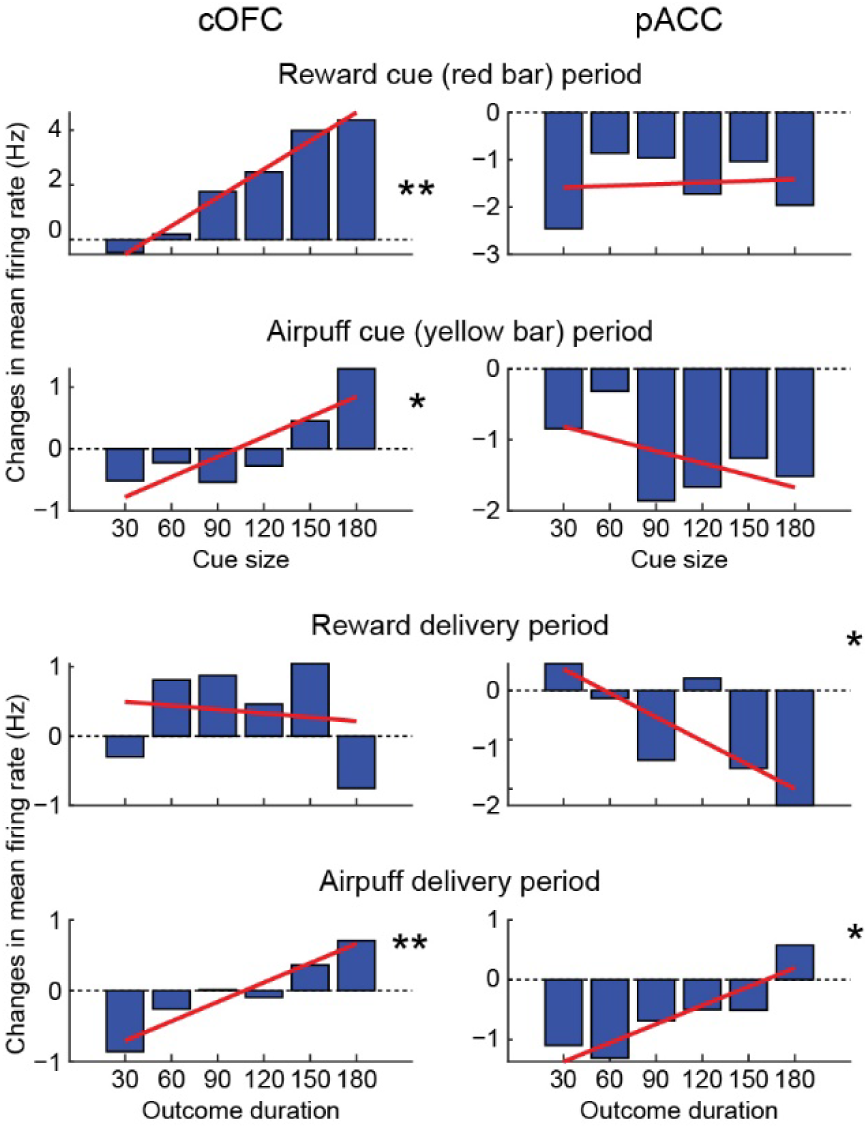
Pavlovian Trials firing rates during cue and outcome periods in the Ap-Av task. A linear trend line (in red) was fitted over these bar values using linear regression to observe the overall trend across different groups. This approach allowed for the visualization of both individual group differences and the overall trend in changes in neuronal activity. The x-coordinates for the bars represented the indices of different sizes of reward/punishment cues or their respective outcomes, whereas the y-axis displayed the mean changes in firing rates between two time intervals. The linear regression was used to model this relationship, and the fitted line provided a visual representation of the trend. The analysis calculated the standard error of the slope, the t-statistic, and the corresponding two-tailed p-value to assess the significance of the trend. *p < 0.05; **p < 0.001.

For the cOFC, we found significant positive trends for both the Reward Cue (t(4) = 14.21, p < 0.0001) and the Airpuff Cue (t(4) = 3.19, p = 0.033), indicating substantial changes in firing rates post-event. By contrast, the pACC trends for both the Reward Cue (t(4) = .2, p = 0.851) and the Airpuff Cue (t(4) = −1.35, p = 0.250) did not reach significance, as determined by the linear regression analysis mentioned above. During the Reward Delivery event, the cOFC did not exhibit a significant trend (t(4) = −0.29, p = 0.784), but the pACC exhibited a marginally significant effect (t(4) = −2.55, p = 0.063). For the Airpuff Delivery event, both the cOFC (t(4) = 6.33, p = 0.003) and the pACC (t(4) = 3.94, p = 0.017) showed significant positive trends, again indicating substantial changes in firing rates post-event.

### cOFC EMS alters approach-avoidance decision-making

We performed 93 cOFC EMS sessions in both monkeys, and identified 29 sites at which EMS induced a significant behavior change (14 sites total for monkey P and 15 sites total for monkey D; Fisher’s exact test, p < 0.05). We used 1-1.5 s long trains of biphasic pulses coincident with the choice cue period onset, with either high (150-200 μΑ) or low (5-15 μA) currents (Fig. 1; green dotted rectangles). Each site was stimulated during the middle block of the Ap-Av EMS task. To perform EMS in a session, we required that choice behavior on the Ap-Av task be stable across at least two consecutive days. Panels a and b of Fig. 7 illustrate the results from a single stimulation session applied in the cOFC in which non-significant change in the Ap-Av choices was observed. Of the 29 significant sites, 23 (79.31%) demonstrated a decrease in Ap and an increase in Av, 4 (13.79%) elicited decreases in both Ap and Av, and 2 (6.89%) elicited increases in both Ap and Av during the EMS block. During the sessions in which we observed statistically significant EMS effects (Fig. 7c-f), the effects (increases in avoidance/approach choices) were generally more pronounced at the beginning of a block and gradually faded out as the block progressed (Fig. 7g). To analyze this temporal pattern, we divided each block of trials into five equal segments and calculated the percentage of Ap and Av trials within each segment, excluding error trials. Relative change was determined for each segment by comparing the corresponding segment of the second block (Stim-on block), during which EMS was delivered, with the matching segment of the first block (Stim-off block), during which no EMS was applied. Nevertheless, there remained a statistically significant increase in avoidance throughout almost the entirety of a block (80%) relative to the sessions without a EMS effect (Fig. 7g). There was a statistically significant decrease in approach trials in 60% of the block’s trials, again compared to sessions without a significant EMS effect. Thus, overall, the EMS of the cOFC increased avoidance behavior. In this analysis (Fig. 7a-f), Fisher’s exact test was used to evaluate whether there was a statistically significant difference in choice behavior between two blocks of trials. For the entire data set, consisting of Ap and Av choices, the choices were divided into those in Block 1 (Stim-off block) and those in Block 2 (Stim-on block). The overall difference in choice behavior between the two blocks was assessed using Fisher’s exact test, which compared the total counts of Ap and Av choices to determine whether there was a significant association between the block and the choice behavior. Separate Fisher’s exact tests were then conducted for the Ap and for Av choices to identify whether there were separate increases in these behaviors. For Ap choice, we compared the proportion of Ap choices between Block 1 and Block 2. A statistically significant increase in Ap choices in Block 2 compared to Block 1 would indicate an increase in Ap behavior; a significant decrease would indicate a reduction. Similar analyses were done for Av behavior to dicate whether there was an increase or decrease in Av behavior in Block 2 compared to Block 1. These findings suggest that the cOFC plays a critical role in modulating avoidance behavior by influencing decision-making processes involving cost-benefit evaluations. Specifically, the cOFC appears to encode a negative bias in valuation, enhancing the weighting of aversive outcomes over rewards. The increase in avoidance behavior induced by EMS of the cOFC indicates that this region contributes to the assessment of potential risks, promoting actions that favor avoidance when potential punishments are present. This aligns with the proposed function of the cOFC in integrating aversive and reward-related information to guide adaptive behavior in situations involving Ap-Av conflicts.

**Figure 7.**
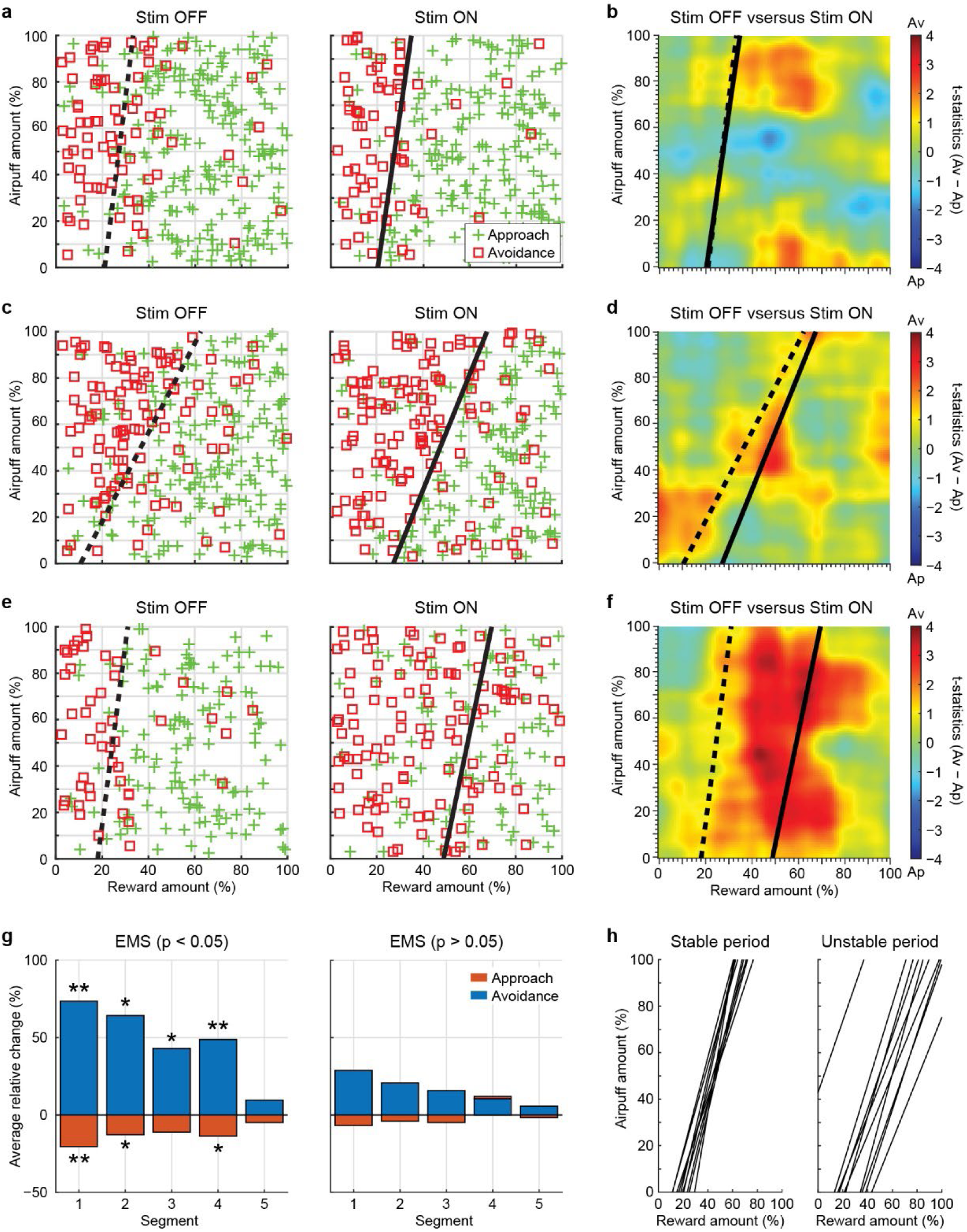
Effects of EMS on decision-making. **a,** A sample EMS session, demonstrating stable behavior (non-significant increase/decrease of approach/avoidance behavior) in the Stim OFF block and Stim ON block during high current EMS (150-200 μΑ). **b,** Matrix plot of t-scores during high current EMS. **c,** EMS session demonstrating a small effect but a significant increase in avoidance behavior during EMS with high currents. **d,** Matrix plot of t-scores during high current EMS. **e-g,** EMS session demonstrating significant effects between the blocks during low current EMS (5-15 μΑ, **e**), and matrix plot of t-scores during low current EMS, between the Stim OFF and Stim ON block (**f**). For panel **g**), each block of trials was divided into five equal segments, and the percentage of Ap and Av trials within each segment was calculated. Relative change of approach and avoidance behavior during sessions where EMS had a statistically significant effect (Fisher’s exact test, p < 0.05, left panel, 29 sessions) and sessions that did not have a statistically significant effect (right panel, 64 sessions; see Methods). **h,** Decision boundaries from 10 consecutive testing sessions in which the monkeys exhibited stable behavior (left panel) and those in which the monkeys exhibited non-stable behavior (Fisher’s exact test, p < 0.05). See methods for details on the construction and analysis of decision matrices.

We performed a basic event-related analysis (Fig. S5a). The results from the EMS blocks largely mirrored those from the standard Ap-Av sessions, showing a predominance of the cOFC across most task events, except for reward delivery events in both Ap and Av choices, where the cOFC appeared more active than the pACC. This pattern contrasts with the standard sessions, in which the airpuff delivery was the event forwhich the pACC appeared more active. Notably, two of the three reward delivery events—task avoidance reward onset (Av rew ON) and Pavlovian reward delivery (Red bar REW)—along with the highlight around cross for choosing Ap (Ap highlight) were the only events that showed a statistically significant difference in spike counts in favor of the cOFC over the pACC (Av rew ON: chi-square test = 6.54, p = 0.01; Red bar REW: chi-square test = 14.22, p = 0.0002; Ap highlight: chi-square test = 5.76, p = 0.016). There was also an increase in the percentage of neurons showing statistically significant activity during the microstimulated cue events (Choice cue Onset, Red bar Onset, Yellow bar Onset), though this result could be partially confounded by EMS artifacts. Thus, following the cOFC EMS, a higher proportion of cOFC neurons exhibited significant activity in response to reward, suggesting a potentially greater involvement or sensitivity of the cOFC towards positive outcomes compared to the pACC. For the Pavlovian trials’ firing patterns during EMS (Fig. S5b), there was a significant negative parametric modulation of the firing rates proportional to the size of the offered cue/outcome (t(4) = −4.48, p = 0.011). This finding suggests that cOFC EMS may alter how the brain processes reward-related information, potentially reducing sensitivity to larger rewards. Such a shift could explain the increased avoidance behavior observed following both low and high current EMS.

In addition to the electrophysiological data, we made a series of physiological measurements, including two autonomic (heart rate variability, pupil diameter) and two somatic (lick activity and reaction times) factors (Fig. 8). Remarkably, we observed that all the physiological metrics were significantly correlated with the behavior of the animals, as well as across each other.

**Figure 8.**
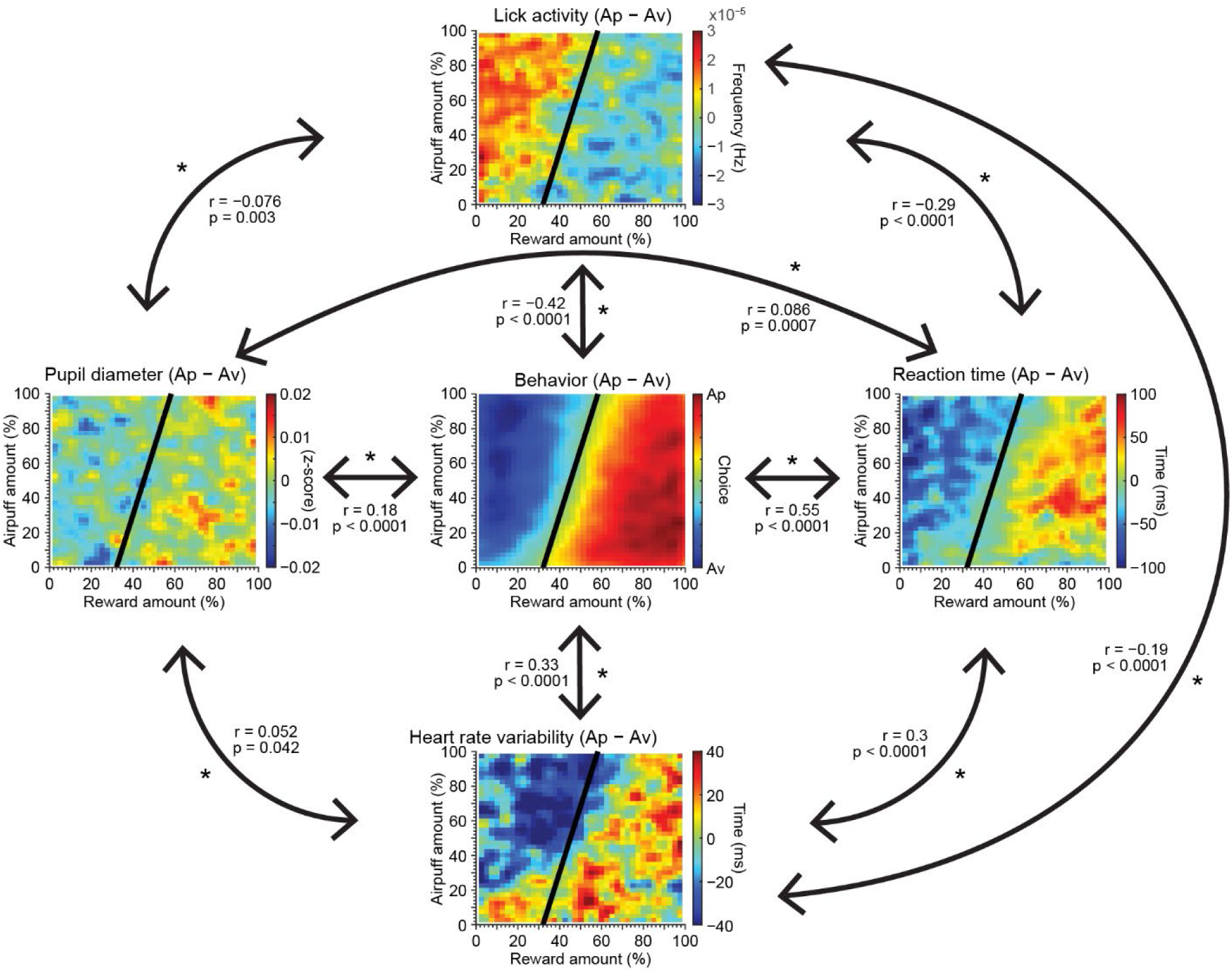
Correlations between physiological metrics and behavioral responses during Ap-Av. The metrics displayed include lick activity, reaction time, heart rate variability, and pupil diameter, averaged from cue onset to outcome onset. Each subplot shows these metrics against reward amount (%) and airpuff amount (%). Significant correlations are highlighted with arrows and correlation values. The central panel consolidates these findings, depicting behavior as a function of reward amount (%) and airpuff amount (%), and emphasizing the collective impact of these physiological measures on the decision-making process in the Ap-Av task. The decision boundary in the physiological data plots is identical to the decision boundary during the respective behavior, for illustration purposes.

For all four, we calculated the mean activity separately for Ap and Av trials, including the period from the choice cue onset to the right before the outcome period onset, approximately 3.7 s, and then we took the difference between Ap and Av for each metric. The matrices displayed around the ‘Behavior’ matrix in Fig. 8 correspond to differences in Reaction Time, Heart Rate Variability, Pupil Diameter, and Lick Activity, plotted as the difference between Ap and Av choices.

For the Pavlovian trials, which had a shorter duration (∼3 s) due to the absence of the decision/choice component (peripheral targets), we also calculated the average activity for the metrics other than reaction time from the single cue (red or yellow bar) presentation until right before the outcome period. We observed distinct patterns of variation with outcome size/duration in lick activity, heart rate variability (HRV), and pupil diameter across reward and airpuff trials (Fig. S6). Lick activity was higher in reward trials compared to airpuff trials and decreased at the smallest reward size. HRV followed a U-shaped pattern during both reward trials and airpuff trials. Pupil diameter changes were more pronounced during airpuff trials, with a notable decrease in diameter as airpuff sizes increased, whereas pupil dilation varied little with reward size during reward anticipation. For lick activity only, we included 5 s before the cue period (which includes the 1-s fixation period and most of the inter-trial intervals) and the outcome period until the offset of the reward delivery event for Ap and Av trials as well as Pavlovian trials, as we wanted to learn how the lick activity changed throughout the course of a full trial (Fig. S7). In Choice trials, lick activity peaked at reward delivery, with small variations at earlier trial events. The same pattern was visible in Airpuff Pavlovian trials, but in Reward Pavlovian trials licking peaked most strongly towards the end of the fixation period.

HRV during choice trials was significantly correlated (Pearson r with observed behavior (r = 0.33, p < 0.0001), reaction times (r = 0.3, p < 0.0001), pupil diameter (r = 0.052, p = 0.042), and negatively correlated with lick activity (r = −0.19, p < 0.0001). Lick activity was negatively correlated with observed behavior (r = −0.42, p < 0.0001), pupil diameter (r = −0.076, p = 0.003), and reaction times (r = −0.29, p < 0.0001). Pupil diameter (r = 0.18, p < 0.0001) and reaction time (r = 0.55, p < 0.0001) were both significantly correlated with observed behavior. Analyses of the lick activity during the entire trial duration for both choice trials exhibited an equal increase in licking activity for both trial types (Ap or Av) before the monkeys received information about the offer (Fig. S7a). Once the offer was presented, a slightly elevated activity occurred for the trials ending in approach (green color), continuing up until the outcome periods. During these outcome periods, elevated licking activity occurred at different time points corresponding to the reward delivery times. During the Pavlovian trials (Fig. S7b), there was a remarkable elevation in licking activity during the fixation point for trials in which the monkey would freely receive a reward (green color). This suggests that the monkeys anticipated the Pavlovian reward trials, which were consistently presented 7-8 trials after the Pavlovian punishment trials.

### Modeling EMS effects in reinforcement learning agents

To explore the dynamics of adaptive strategies under the influence of EMS, we employed linear advantage actor-critic (A2C) models, a subclass within the broader spectrum of reinforcement learning algorithms (Mnih et al., 2016). Our model aimed to capture the general behavior patterns observed across both monkeys and all sessions in the Ap-Av task, rather than fitting specific individual sessions.

In the Ap-Av task, the monkeys were considered to use both potential rewards (x1) and risks (x2) in making their decision, as described by Levy & Glimcher (2010). Here, the decision metric d is calculated as:

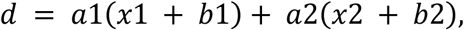

where a1 and a2 represent the weights the agent assigns to each observation, signifying their relative importance, and b1 and b2 are biases that adjust the baseline values of the observations. The decision to approach or to avoid is not made directly from d, but rather through converting d into a probability via the logistic function (McFadden, 1972):

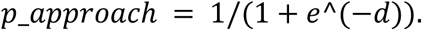

This probabilistic approach allows for nuanced decision-making, encapsulating the agent’s tendency towards either approaching or avoiding based on its calculated confidence level. Our experimental findings for the Ap-Av task indicated that EMS predominantly influenced the monkeys by skewing decisions towards avoidance. To simulate the effect of the EMS in our model, we introduced a parameter γ, modifying the decision metric to:

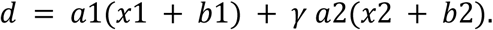

The γ parameter allows us to modulate the effect of the aversive stimulus (x2) on the decision-making process. When γ > 1, it amplifies the weight of the aversive stimulus, simulating the effect of EMS in increasing avoidance behavior.

In Fig. 9, we illustrate the effect of cOFC EMS on the A2C model and how varying levels of EMS influence the agent’s decision-making capabilities. The model’s baseline decision boundary without EMS is shown with a logistic regression line (black, Fig. 9a,c). The adjusted decision boundary with EMS (γ = 1.25), shown by a grey logistic regression line in Fig. 9b and c, indicates the EMS’s effect on the model’s evaluation of risks and rewards. The strategic shift in bias induced by the cOFC EMS is illustrated by the contrast between these two decision boundaries, emphasizing the flexibility and adaptability of the model in response to external stimuli (Fig. 9c). The parameter gamma (γ) directly correlates with EMS intensity because it mathematically represents how much the brain’s risk processing is amplified during stimulation. Think of γ as a dial that adjusts the “volume” of risk perception—when γ = 1, there’s no stimulation, and risk is processed normally. As we increase γ above 1 (e.g., to 1.25 or 1.50), it is analogous to increasing the stimulation current, which makes the brain’s risk-detection circuits more sensitive. This results in a systematic shift in the decision boundary: higher stimulation currents (represented by higher γ values) lead to stronger avoidance behaviors, even when the objective risk levels remain the same. It’s similar to turning up the brightness on a display, making everything appear more intense—higher γ values make risks appear more threatening to the decision-making system. Moreover, incremental changes in gamma values (from 1.10 to 1.50) systematically alter the agent’s propensity to engage in avoidance behavior (Fig. 9d). The gradation of EMS intensity is directly correlated with the shift in the decision boundary, reflecting an enhanced sensitivity to risks over potential rewards. This effect underlines the modulation of the monkeys’ assessments of aversive stimuli versus rewards by the EMS, thereby shaping the overall strategy in the Ap-Av task.

**Figure 9:**
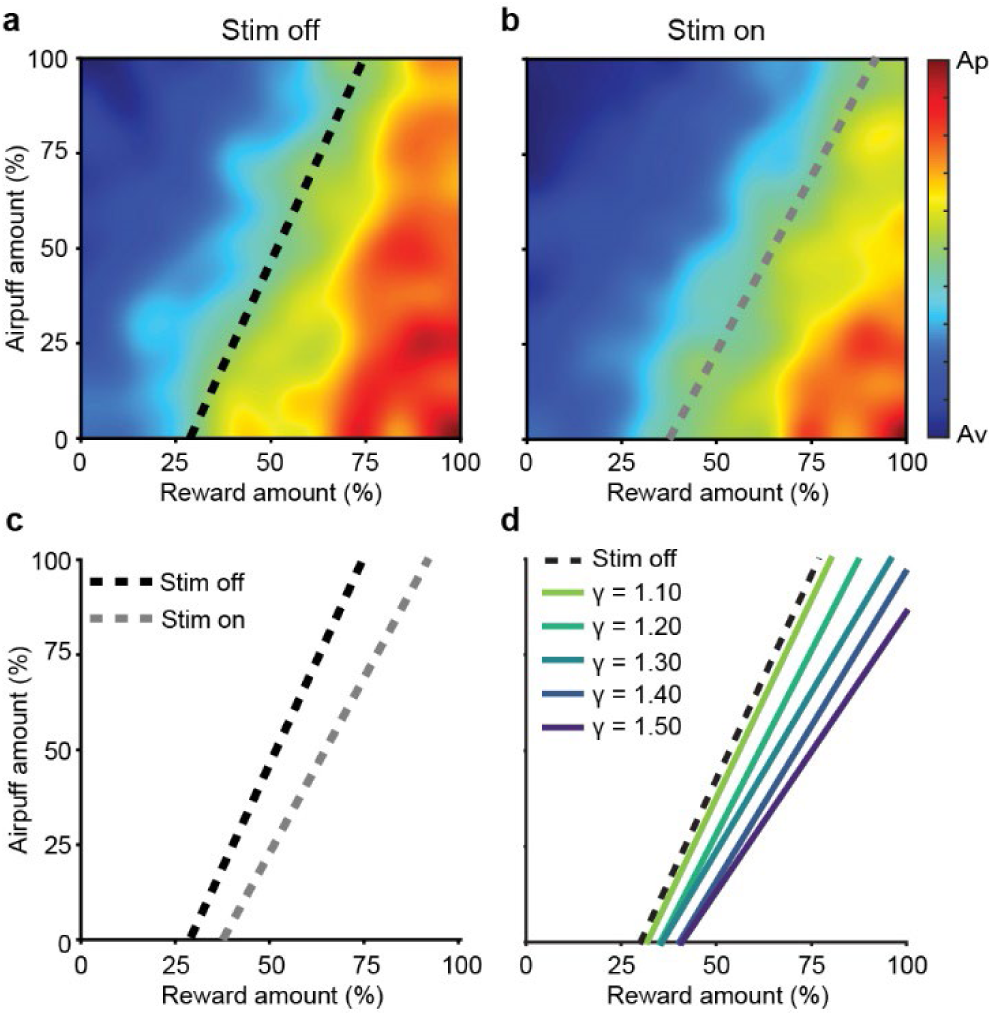
Comparative analysis of the A2C model’s behavior under the influence of EMS. **a,** Baseline decision boundary without EMS, depicted by a dotted line. **b,** Adjusted decision boundary when EMS was applied (γ = 1.25), with the solid line illustrating the shift in strategy. **c,** Direct comparison of these decision boundaries, highlighting the significant impact of the EMS on the model’s Ap-Av decisions. This depiction underscores the model’s adaptability and the profound effect of external stimuli on decision-making processes. **d,** The effect of varying levels of EMS on the decision boundary. This figure demonstrates how different EMS intensities (ranging from 1.10 to 1.50) influence the decision-making process in an approach-avoidance task. The y-axis represents the strength of the airpuff, a proxy for the aversive stimulus, whereas the x-axis indicates the reward amount. Each line corresponds to a different level of EMS, illustrating a gradual shift in the decision boundary. This shift signifies the agent’s increased inclination towards avoidance with higher EMS levels, elucidating the potent effect of the EMS on modulating the balance between risk and reward in decision-making.

Our model thus successfully captured the general trends observed in monkey behavior across multiple sessions, demonstrating increased avoidance behavior under EMS conditions. This reinforcement learning approach provides a computational framework for studying the neural mechanisms underlying decision-making in such Ap-Av conflicts and the potential modulatory effects of cOFC on these processes.

## Discussion

Our findings in macaque monkeys point to four main conclusions. First, there are significant differences in the neural activity of well-isolated units recorded in the cOFC and pACC during the performance of both the Ap-Av conflict decision-making and Pavlovian trials. Units in the cOFC demonstrated a balanced response pattern with similar percentages of excitatory and inhibitory responses, particularly in response to airpuff and reward cues. This balanced activity suggests that the cOFC is likely involved in integrating both positive and negative stimuli, a capacity essential for successful decision-making and emotional regulation. The equal distribution of excitatory and inhibitory responses supports evidence that the cOFC can dynamically adjust its output to respond appropriately to varying motivational contexts, thereby maintaining flexible and adaptive states (Wallis, 2007; Rolls, 2004). By contrast, units in the pACC exhibited a predominance of inhibitory responses: a larger proportion of pACC neurons decreased their firing rates more in response to negative than they had excitatory responses. This might suggest that the pACC is important in modulating or suppressing neural activity, potentially refining or filtering incoming information and contributing to selective engagement in specific task aspects (Etkin et al., 2011). The high level of unresponsiveness in these regions (c.f., Amemori and Graybiel, 2012) suggests that they likely are engaged in processes outside the immediate scope of our measurements, including network effects (Amemori et al., 2024) or that they have a more selective role in responding to certain types of stimuli (Shackman et al., 2011).

Second, patterns of response during the cue period across both choice and Pavlovian trials in the cOFC and pACC were stable, suggesting that these regions could share a function in inducing or reflecting this stabilization. The analyses during the Pavlovian trials helped to delineate the differential involvement of cOFC and pACC neurons in response to single rewarding and aversive cues (Fig. 6). In the cOFC, both cue events show a significant positive trend, indicating a substantial increase in firing rates post-event. The pACC, however, shows a significant positive trend during both *outcome* events, suggesting increased neuronal activity in response to these events. Other events either showed non-significant trends or trends that were not strong enough to reach conventional levels of statistical significance. We conclude that the cOFC is significantly activated by both positive and negative stimuli, indicating its crucial role in anticipating rewards and punishments. Conversely, the pACC responds both to positive and negative outcomes, highlighting its involvement in processing potential rewards, threats or losses. These findings suggest that distinct neural circuits are involved in evaluating different aspects of cost-benefit decisions, providing essential insights into how the brain navigates complex decision-making scenarios that entail both positive and negative outcomes. These findings have significant implications for understanding the neural mechanisms underlying mood-related disorders, such as major depressive disorder (MDD). The balanced excitatory responses to salient rewarding and aversive offers in the cOFC that we observed in the monkeys might reflect a neural mechanism that, if conserved in humans, could, when disrupted, contribute to MDD symptoms such as anhedonia or heightened sensitivity to either positive or negative stimuli (Drevets et al., 2008). The predominant inhibitory response in the pACC could indicate a mechanism for regulating values derived from the integration of positive and negative emotional responses and cognitive processes, potentially contributing to the dysregulation seen in MDD (Mayberg, 1997). However, much further detailed analysis is needed.

Thirdly, our findings demonstrated that EMS of the cOFC with both high (150–200 μA) and low (5–15 μA) currents effectively induced avoidance behavior in the subjects, indicative of a pessimistic shift in decision-making. This effect was not only observed behaviorally but was also captured by our reinforcement learning model, as detailed in Fig. 9. The model showed that EMS could be simulated by increasing the gamma (γ) parameter, which amplifies the weighting of aversive outcomes over rewards. This shift in γ resulted in a systematic increase in avoidance behavior, mirroring the behavioral changes observed in the monkeys. Therefore, our analyses suggest a robust causal relationship between cOFC activity and the modulation of decision-making processes, particularly by enhancing the perceived cost in cost-benefit analysis. Moreover, the consistent elicitation of avoidance behavior through EMS underscores the role of the cOFC in encoding a negative bias during valuation processes. This finding aligns with evidence suggesting a “stop function” for this region in inhibiting actions that may lead to negative outcomes (Robbins, 2007; Aron et al., 2014). These results are consistent with previous research indicating the involvement of other parts of the OFC in emotional regulation and decision-making under risk (Bechara et al., 2000; O’Doherty et al., 2001).

Finally, our experiments have uncovered significant correlations between these neuronal activities and different classes of autonomic and somatic activities recorded during Ap-Av conflict decision-making performance. The significant correlation between HRV and observed behavior aligns with previous studies demonstrating that HRV is a reliable indicator of emotional and stress responses, which can influence decision-making (Thayer et al., 2012; Appelhans & Luecken, 2006). The positive correlation between the difference in HRV and the difference in reaction times further supports the notion that autonomic activity supports the performance of cognitively demanding tasks, in this case, saccading to the correct target to implement the monkey’s Ap or Av decision (Thayer & Lane, 2000). Similarly, the correlation between pupil diameter and behavior highlights the importance of arousal and attentional mechanisms. Pupil diameter is a well-established marker of cognitive and emotional processing (Bradley et al., 2008; Nassar et al., 2012). Increased pupil diameter is associated with greater cognitive load and decision uncertainty (Murphy et al., 2014). The negative correlations between the difference in lick activity and both observed behavior and the difference in reaction times during choice trials suggest that licking behavior, potentially indicative of anxiety or arousal, inversely affects decision speed and accuracy. Our data indicate that monkeys lick more during Ap trials with high airpuff and small reward cues compared to Ap trials with large rewards and small airpuff cues, suggesting heightened anticipatory behavior or anxiety in these conditions (Fig. S8). This form of anticipatory anxiety has been observed in many species, including monkeys, highlighting its role in preparing animals for expected stressors and its impact on behavioral responses (Kalin et al., 1991; Kalin et al., 1996). The observed relationships among these physiological metrics further emphasize their interconnected nature. For instance, the significant correlation between HRV and pupil diameter suggests a link between autonomic regulation and attentional mechanisms, corroborating the idea of a shared underlying system influencing both (Sammito & Thielmann, 2020). anticipatory anxiety can lead to increased stress responses and behavioral changes, highlighting the importance of these responses in understanding the pathology of mood disorders and the potential for targeted therapeutic interventions (Folkedal et al., 2012; Gygax et al., 2013). Thus our findings suggest strong ties between neural activities in the cOFC and pACC and both autonomic and somatic components of behavior accompanying these activities. We suggest that these findings could contribute new insight into the brain-body axes that are characteristic of salient motivationally challenging behavioral states.

The primate pACC and cOFC are reciprocally connected (S. Amemori et al., 2021) and send outputs to subcortical structures. Notably, the pACC, along with the subgenual anterior cingulate cortex (sACC) and the cOFC, are limbic cortical regions that receive direct inputs from the amygdala (Ghashghaei et al., 2007). Both the pACC and cOFC have preferential projections to the striosome compartment of the striatum, especially to the anterior striosomes of the caudate nucleus and the adjoining rostral part of the putamen (Eblen and Graybiel, 1995; S Amemori et al., 2020). Here, we conducted EMS experiments for functional mapping of the cOFC and demonstrated its causal involvement in pessimistic decision-making, similar to the function of the pACC. Furthermore, we performed neuronal recording to address the functional differences between the pACC and cOFC. Importantly, we found differential neuronal responses during the Ap-Av decision-making. The responses of the cOFC reflected physiological measurements that were positively correlated with behavioral patterns, emphasizing body-brain synchronization in the cOFC. These results uncovered a functional dichotomy between the pACC and cOFC pathways, which send different negative evaluation signals related to pessimistic value judgments compatible with an anxiety-like state, targeting the striosome compartment of the striatum.

## Methods

### Subjects

Two Macaca mulatta monkeys, one male (P, 15.4 kg) and one female (D, 9.1 kg), were studied in experiments that strictly adhered to the Guide for Care and Use of Laboratory Animals of the United States National Institutes of Health. All experimental procedures were approved by the Committee on Animal Care of the Massachusetts Institute of Technology. Prior to training, both monkeys underwent a habituation process that acclimated them to a seated position in a monkey restraining chair and ensured a proper facemask fit. All surgical procedures were performed under sterile conditions with deep anaesthesia. Postoperative analgesics were administered to the monkeys. Prophylactic antibiotics were injected intramuscularly on the day of the surgery and were continued daily for the subsequent week.

### Task procedures

The initial phase of training aimed to acquaint naive monkeys with the testing environment and basic task structure. Monkeys were first presented with a large central white fixation point occupying half of the screen. The goal was to have them fixate on this point, beginning with short durations, such as 50 ms. Over time and with consistent training, the size of this fixation point was gradually reduced, and the duration of required fixation was increased. This progression continued until monkeys could consistently maintain their gaze for up to 1,000 ms. After establishing consistent fixation behavior, the monkeys were introduced to the red and yellow bars. At this introductory stage, both bars represented positive outcomes. Different sizes of these bars were displayed to match different sized outcomes to emphasize the significance of size in the upcoming tasks and to instill the understanding that ‘size matters’.

As the training progressed, specific associations were assigned. The red bar continued to be associated with a positive outcome and was now consistently linked with a white cross. The yellow bar, which had initially indicated a positive outcome, was now consistently linked with a white square (undergoing a transformation later in the Ap-Av task, when it began to signify a negative outcome). Training continued with the Ap-Ap task, in which both choices led to positive outcomes. During a typical choice trial, the sequence began with the monkey fixating on the central fixation point for up to 1,000 ms. Then, the cue offer was presented, with two abutted red and yellow bars of random sizes, followed by peripheral symbols representing the outcomes linked to the red (cross) and yellow (square) bars, which appeared for 1 s. The monkey was required to fixate on one of these symbols for at least 200 ms, indicating its choice. A successful choice was confirmed with a yellow highlight circle around the selected symbol, whereas a failure to decide within the given time rendered the trial invalid.

Following a correct choice, an interval of 600-800 ms preceded the outcome, which corresponded to the chosen bar’s association. If fixation was broken at any point during the task, a penalty in the form of a 5-s delay was imposed before the initiation of the next trial. Upon mastering the Ap-Ap task, monkeys moved to the Ap-Av task, in which the choices were associated with contrasting outcomes: positive for the red bar and negative for the yellow bar. In the Ap-Av task, the concept changes; instead of choosing between the two bars, the monkeys had to decide whether to approach or to avoid the presented offers by making a saccade to the cross (to approach) or to the square (to avoid). After solidifying their understanding of the Ap-Av task, monkeys tackled the Ap-Av-Ap-Ap format. The Ap-Av (first and third block) and Ap-Ap (second and fourth block) tasks were given in an alternating sequence. Mixed into these primary tasks were the Pavlovian cue trials, which required no decision-making and constituted around 15% of all trials. The trials consisted on bars of certain sizes/dimensions: 30 (bar size: 5.5 mm x 8.5 mm), 60 (bar size: 11 mm x 8.5 mm), 90 (bar size: 16.5 mm x 8.5 mm), 120 (bar size: 22 mm x 8.5 mm), 150 (bar size: 27.5 mm x 8.5 mm), and 180 (bar size: 33 mm x 8.5 mm).

### Recording setup

After behavioral training phase, individualized recording chambers, informed by 3T MRI coordinates, were implanted. These coordinates were derived from T1-weighted (0.5 mm isotropic) and T2-weighted (0.35 mm isotropic) MRI scans, which provided a detailed view of the skull surface. Gray Matter Research fabricated these chambers, which were implanted on both monkeys (P and D) so as to allow access to a significant part of the orbitofrontal cortex and the anterior striatum. Several weeks following the chamber installation, a craniotomy was performed to expose the dura mater. The chamber was sealed using silicone elastomer (Kwik-Sil) applied directly to the dura mater. To pinpoint the exact coordinates for each electrode track, an MRI scan made with the chamber and grid (Feingold et al., 2012) filled with a solution of saline combined with a 5% gadolinium contrast agent. Approximately two weeks, the electode array implant surgery took place. Chambers were fitted with grids that featured an array of openings (40 mm × 30 mm), each with a diameter of 0.48 mm and a center-to-center distance of 1 mm. Probes were secured onto microdrives, so that they could be inserted by screw-controlled adjustments (158 µm per complete turn). Custom-made micromanipulators held all these electrodes in place on the grid. The MRI (T1 and T2-weighted images, 0.5 mm slice thickness) with the chamber and grid infused with saline and the 5% gadolinium contrast agent allowed the coordinates of each electrode track to be determined. Under anesthesia and in sterile conditions, platinum-iridium electrodes (with impedances ranging between 0.8-1.5 MΩ; FHC Inc., Bowdoin, ME) were implanted. For Monkey P, 57 electrodes were implanted: 20 in the pACC and 37 in the cOFC. Monkey D was implanted with 48 electrodes: 15 in the pACC vicinity and 33 in the proximity of the cOFC.

### Recording of physiological activity

To characterize the internal behavioral states of the monkeys during task performance, we measured licking, pupil diameter, pulse, and reaction time. Licking was quantified by summing the absolute values of mouthpiece acceleration in three dimensions, using data from a three-axis accelerometer (SparkFun, MMA8452Q) attached to the mouthpiece that delivered liquid reward. These signals were directly routed to the input of the electrophysiology system (SX Neuralynx) after being attenuated to match the system’s input range (±5 mV). This allowed synchronous recording with neural electrical activity at the same sample rate. Further processing of the signals was performed using MATLAB 2018a, in which relevant features were extracted from the raw data. The three-dimensional licking signal was obtained by summing the absolute values of the signals from the X, Y, and Z dimensions. The combined signal was low-pass filtered at 10 Hz to remove high-frequency noise. The filtered signal was then downsampled from 32 kHz to 1 kHz to reduce data size and to facilitate further analysis. The processed licking signals were saved for subsequent analysis.

To generate the graphs shown in Fig. S7, the licking activity for each trial was first interpolated to match the average trial length of each trial type in each session with MATLAB’s ‘interp1’ function. This standardization was done to ensure consistency across the same type of trials within each session. At each time point, an average was created across all trials. The trial average time course was then interpolated for each session to match the average trial length across all sessions. The interpolated waveforms where then averaged across all sessions. Finally, error bars were calculated to represent variability as the standard error of the mean across sessions.

Pupil diameter was measured using an infrared eye-tracking system (SR Research, EyeLink 1000) designed for the visually guided task, with signals also routed to the electrophysiology system post-attenuation. Regions where the monkey exhibited sleepiness and generated spurious “blink” signals were identified based on the density and abnormal duration of these signals. The pupil diameter was set to a continuous blink-level value in these regions, making them appear similar to periods when the monkey’s eyes were fully closed. These cleaned data were low-pass filtered and downsampled by a factor of 32. The resulting file was used to calculate the average and median pupil diameter from cue onset to outcome onset for each trial, with values for choice and Pavlovian trials saved separately in a .mat file.

Pulse measurements were obtained from an ear-clipped oximeter (SparkFun, SEN-11574), processed similarly to licking and pupil diameter signals, and downsampled to 1 kHz. HRV metrics were derived by detecting peaks in the z-score normalized oximeter signal, calculating intervals between peaks, and computing the standard deviation of these intervals (RRstd) within defined target-period task windows.

Reaction time was defined as the time needed between choice cue offset and the choice to saccade on one of the two peripheral targets, cross or square symbol, for, respectively, choosing Ap or Av. These were recorded from our electrophysiology system as well as from the behavioral task system (NIMH MonkeyLogic) controlling event sequences (Hwang et al., 2019).

We calculated the difference in average reaction times between approach and avoidance choice trials across all session types. Reaction times were binned based on reward and airpuff levels, with each bin covering 5% increments for both reward and airpuff amounts. For each session, average reaction times for approach and avoidance trials were separately computed by summing and counting within each bin. The difference was then taken between the Ap reaction time and the Av reaction time. To highlight patterns across sessions, these differences were averaged and plotted as heatmaps. We smoothed the reaction time difference data by applying a Gaussian filter using MATLAB’s ‘fspecial’ function to create the filter kernel, and then convolved it with the data using ‘imfilter’, which helped in reducing noise and improving the clarity of the heatmap visualization. A decision boundary derived from the econometric model (see below) was overlaid to indicate regions of behavioral transition between approach and avoidance decisions. The color scale represents the difference in reaction times (approach minus avoidance). The same procedures were applied to all physiological differential Ap-Av matrices.

The relationship between the physiological metrics was further investigated by calculating the Pearson correlation coefficients between pairs of difference variables: pupil size difference, reaction time difference, decision difference, HRV difference, and licking behavior difference. Prior to the correlation analysis, we performed data preprocessing to ensure the validity and reliability of the results. Specifically, we flattened any matrix-form data into one-dimensional vectors to standardize the data structure across all variables. We then removed any observations containing NaN (Not-a-Number) values or zeros in any of the variables, as these could represent missing or non-informative data that might skew the correlation results. This ensured that all data vectors remained aligned and of the same length. The Pearson correlation coefficient was then calculated for each variable pair. Alongside the correlation coefficients, we computed the associated p-values to assess the statistical significance of the observed correlations.

### Econometric modeling

To calculate the internal variables or parameters and to understand the decision-making processes of monkeys, we employed an econometric model to approximate the monkeys’ choice behavior. This model is fundamentally based on the assumption inherent in discrete choice models, where the choice alternatives must be mutually exclusive—choosing one option precludes selecting another—and exhaustive, meaning all possible choices are included and are finite in number. We used the conditional logit model, the most common and widely adopted model for analyzing discrete choice behavior.

Three axioms were essential for the application of this model (Amemori et al., 2012): (1) the subject is a utility-maximizing decision-maker; (2) utilities can be represented as the linear sum of a representative term and an error term, i.e., U_+_ + e_+_ for one choice and U_□_+ e_□_ for the other; and (3) the error term, represented as e_j_ (j = + or □), is independently and identically distributed. Within the context of the sampled monkeys’ decisions, we can infer these subjective values in reverse. If there are two options (denoted as ‘+’ and ‘□’ targets), the probability of choosing ‘+’ targets can be expressed as p_+_ =1/(1+exp(-(U_+_ - U_□_))), where U_+_ and U_□_ represent the representative utilities of each option.

### Stepwise regression

We performed multiple regression analyses to examine the patterns of neuronal activity during the cue period in the Ap-Av task across all session types. To identify an optimal set of variables that linearly parameterized neuronal responses, we used a stepwise regression approach through Matlab’s “stepwisefit” function (Mathworks, Natick, MA). This method iteratively adds or removes variables from a linear model based on their statistical significance, as determined by a series of F-tests. It starts with a preliminary model and assesses the impact of including or excluding each variable on the model’s explanatory power, using the p-value of the F-statistic as a measure for comparing models with or against each additional variable. The threshold for determining a variable’s significance was a p-value less than 0.05.

In our analysis of the Ap-Av task, the selected variables included: Rew (reward value indicated by a red bar), Ave (airpuff duration indicated by a yellow bar), Eutil (expected utility in the Ap-Av scenario), Cho (binary indicator for approach [1] or avoidance [0] choices), Cho*Rew (interaction between choice and reward), Cho*Ave (interaction between choice and airpuff duration), Conf (presence of decisional conflict), and RT (reaction time). These variables were chosen for their potential to elucidate the underlying factors influencing neuronal activity patterns associated with decision-making in these tasks (Fig. 4).

### Pavlovian trials

To assess changes in firing activity across different levels of reward and punishment in the cOFC and pACC, we computed the mean firing rates for 1 s before and after the cue and outcome events and then calculated the difference between these means (Fig. 6). To evaluate the significance of observed trends, linear regression was performed on these mean differences using MATLAB’s ‘fitlm’ function. Each regression involved fitting a first-degree polynomial (a line) to estimate the slope. The fitlm function provided the t-statistic for the slope, calculated as the slope divided by its standard error, which accounts for the variability of the fit. The significance of the slope was assessed directly through the two-tailed p-value provided by the fitlm output. This p-value was used to determine the statistical significance of the trend across different reward and punishment levels.

### Electrical microstimulation

For stimulation experiments, monopolar stimulation was applied. The stimulation train consisted of 200-μs pulses delivered at 200 Hz. Each pulse was biphasic and balanced, with the cathodal pulse leading the anodal pulse. Task events were also sent to a separate PC running MATLAB (MathWorks) to control the EMS generated by the stimulator (Master-8, A.M.P.I.) and isolator (A365, WPI).

To quantify the effects of EMS, we compared the differences between decision matrices of the Stim-off block and the Stim-on block. Decision matrices were constructed by convolving the choice data with a 30-by-30-point square-smoothing window and aggregating each choice datum at each point in a 100-by-100 grid. We then calculated t-statistics to measure the difference in avoidance and approach frequencies between the Stim-off and Stim-on blocks. Positive t-statistics (Fig. 7b,d,f; shown in red) indicate regions where EMS resulted in a relatively larger increase in avoidance behavior compared to approach behavior. Negative t-statistics (Fig. 7b,d,f; shown in blue) indicate regions where EMS resulted in a relatively larger increase in approach behavior. The Fig. 7 plot highlights specific conditions in which EMS significantly influenced decision-making behavior, either by increasing avoidance, increasing approach, or having minimal effect.

For the analysis presented in Fig. 7g, each block of trials was divided into five equal segments (parts), and the percentage of Ap and Av trials within each segment was calculated, excluding error trials. Relative change was determined for each segment by comparing the corresponding segment of the second block (Stim-on block), during which EMS was delivered, with the matching segment of the first block (Stim-off block), during which no EMS was applied. These relative changes were aggregated across all sessions into two distinct groups: ‘EMS-significant’ and ‘EMS-non-significant’ sessions. For each segment, independent two-sample t-tests were conducted to compare the average relative changes in Ap and Av behaviors between the EMS-significant and EMS-non-significant groups. Statistical significance was determined at p < 0.05 (*), and p < 0.001 (**).This analysis enabled the assessment of whether EMS selectively modulated Ap and Av behaviors across different segments of the task by comparing the proportional changes between the two experimental conditions across all sessions. Significant and not-significant effects were distinguished using Fisher’s test (p < 0.05), to compare numbers of Ap and Av trials across blocks.

### Computational model

We implemented an OpenAI Gym environment to simulate the Ap-Av task. This environment provides a two-dimensional observation space representing reward (x1) and risk (x2) values, both normalized between 0 and 1. The action space is discrete with two possible actions: approach (1) or avoid (0). At each step, a new random observation (x1, x2) is generated. If the agent chooses to approach, it receives a reward equal to the benefit minus the cost. If the agent chooses to avoid, it receives no reward. The episode does not terminate, allowing for continuous interaction.

Our A2C model consists of two main components: an Actor (Policy Network) and a Critic (Value Network). The Actor takes a 2-dimensional observation as input and outputs the probability of approaching. The Critic estimates the state value based on the same input. We trained our model using a custom implementation of the proximal policy optimization algorithm. The training spanned 1,500 epochs, with each epoch consisting of 100 steps. We capped individual trajectories at 100 steps to maintain focused learning experiences. A discount factor (γ) of 0.99 was employed to appropriately value future rewards, and we utilized generalized advantage estimation with λ = 0.95 to balance bias and variance in advantage calculations. To promote exploration, we incorporated an entropy coefficient (β) of 0.2, which could be fine-tuned via command-line arguments.

We chose the Adam optimizer for its adaptive learning rate capabilities, with initial learning rates set at 1e−3 for both the policy and value networks. To prevent explosive gradients, we implemented optional gradient clipping with a configurable maximum norm. The training progress was logged every 20 epochs, with model checkpoints saved at the same interval. We tracked average trajectory rewards and lengths to monitor the agent’s performance over time.

### Histology and imaging

The monkeys were deeply anesthetized with sodium pentobarbital and perfused transcardially with 0.9% saline followed by 4% (wt/vol) paraformaldehyde in 0.1 M NaKPO_4_ buffer (PB). Brains were blocked and stored in 25% glycerol (Sigma-Aldrich, G5516) with 0.1% sodium azide (MP Biomedicals, 0210289190) in 0.1 M PB at 4 °C until sectioning. The brains were then frozen in dry ice, and coronal sections were cut at 40 μm thickness on a sliding microtome. Sections were stored in 0.1% sodium azide in 0.1 M PB until use.

For immunofluorescent staining, sections were rinsed three times for 5 min each in 0.01 M NaKPO_4_ buffer with sodium/potassium saline containing 0.2% Triton X-100 (Sigma-Aldrich, T8787; PBS-Tx), and then incubated in blocking solution consisting of 10% normal goat serum in PBS-Tx for 1 hr. The sections were subsequently incubated with primary antibody solution containing rabbit anti-GFAP [1:500] (DAKO, Z0334) in blocking solution for 24 hr at 4 °C. After primary antibody incubation, the sections were rinsed three times for 5 min in PBS-Tx and then incubated for 2 hr in secondary antibody solution containing goat anti-rabbit Alexa Fluor 647 [1:300] (Life Technologies, A21245) in blocking solution at room temperature. Following three additional 5 min rinses in 0.1 M PB, sections were counterstained with DAPI [1:1000] (Life Technologies, 62248) in 0.1 M PB for 2 min, then rinsed in PB for three 5 min intervals, mounted onto glass slides, and coverslipped with ProLong Gold antifade reagent (Life Technologies, P36930).

Images were captured using an AxioZoom V16 microscope (Zeiss) and exported in JPEG format.

## Supplementary Figures

**Figure S1.**
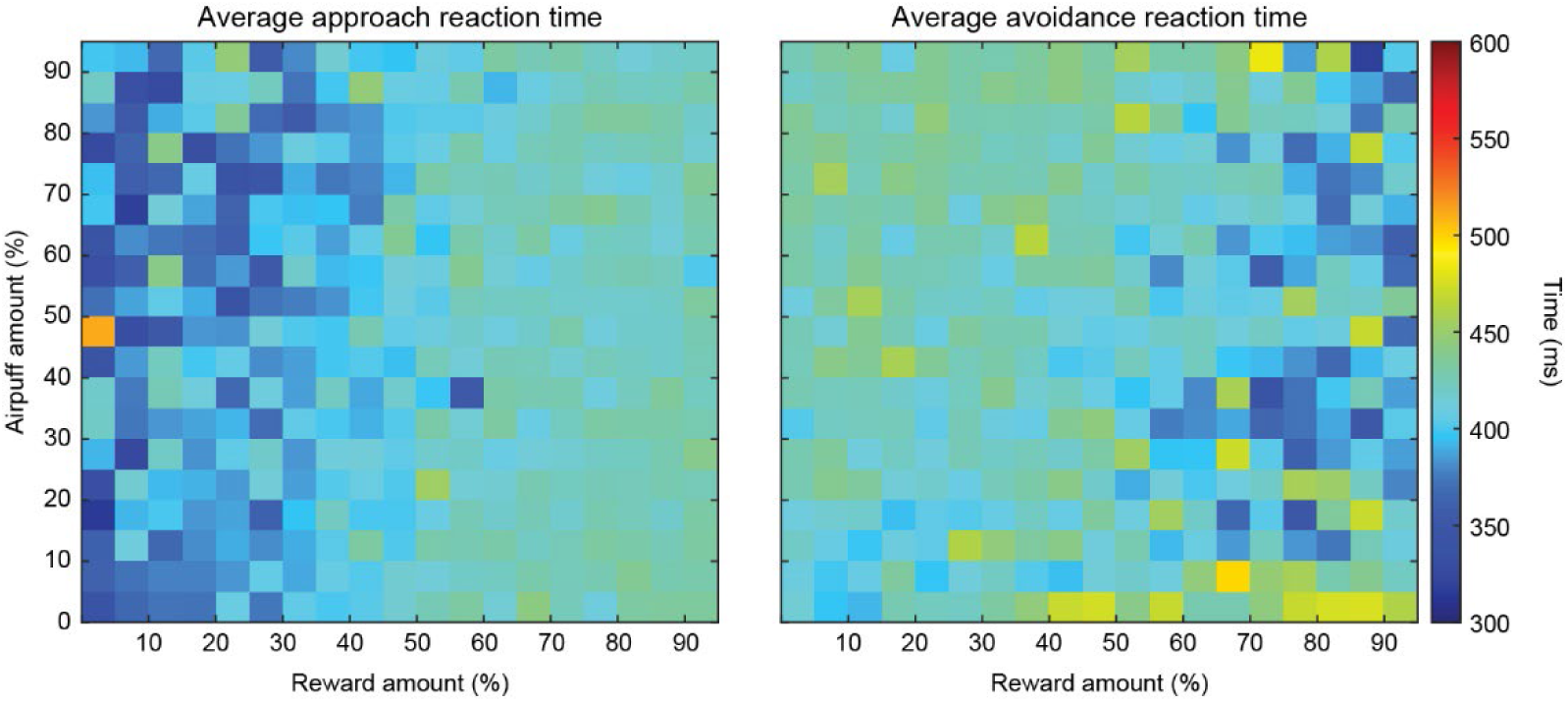
Reaction times during Approach and Avoidance trials across varying reward and airpuff amounts. Heatmaps displaying the average reaction times for approach (left) and avoidance (right) trials. The horizontal axis represents the reward amount (0% to 100%); the vertical axis represents the airpuff amount (0% to 100%). Reaction times are color-coded, with cooler colors (blue) indicating faster responses and warmer colors (yellow/red) indicating slower responses. Monkeys exhibited slower reaction times when approaching better options (high reward, low airpuff) and when rejecting worse options (high airpuff, low reward). This pattern suggests more deliberate decision-making in both cases, wherein animals take longer to approach favorable options and hesitate to reject highly aversive ones.

**Figure S2.**
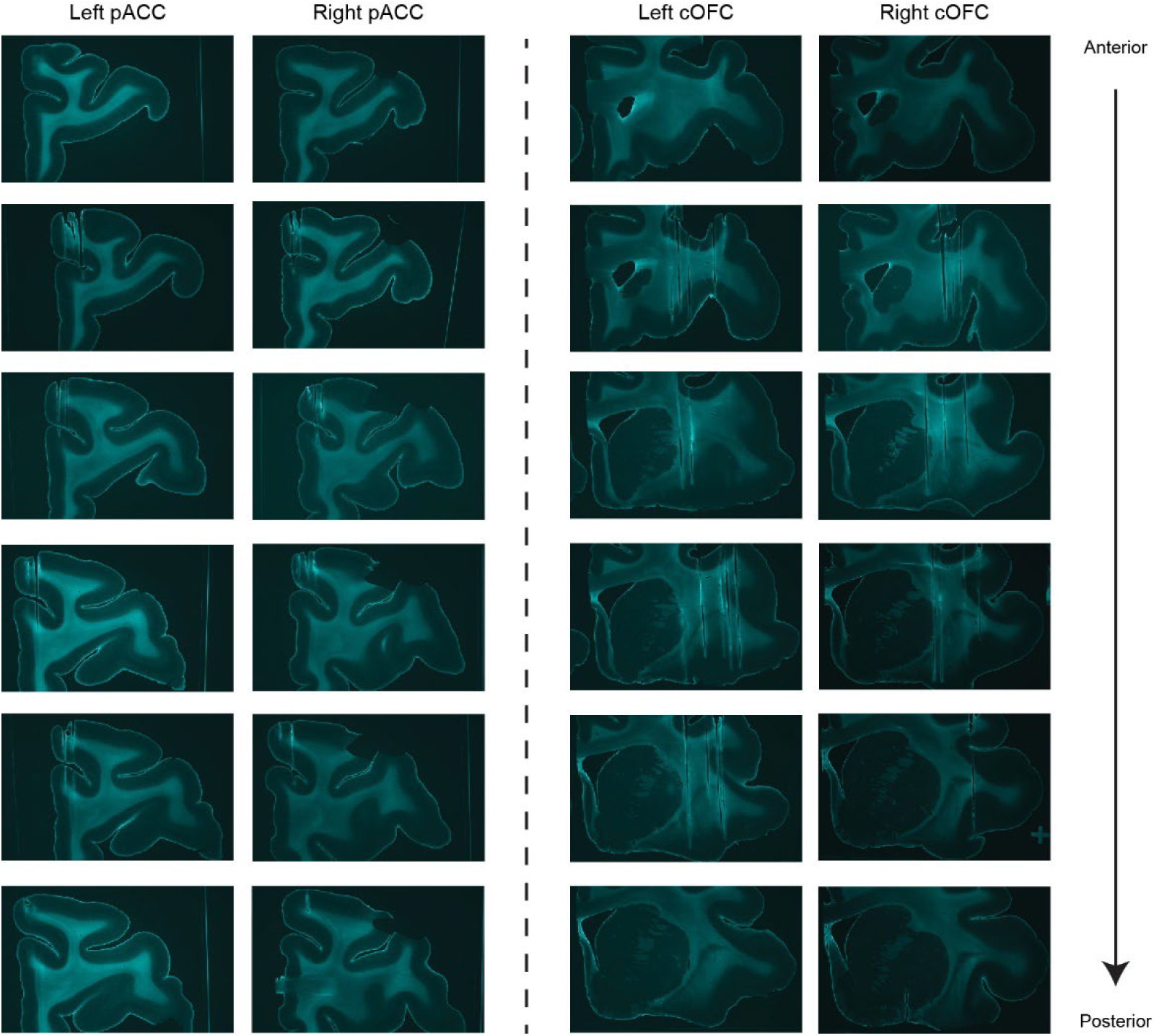
Histological verification of electrodes placements in monkey. **D.** Coronal brain sections from monkey D, arranged from anterior to posterior, illustrating electrode tracks in the pACC and the cOFC. For each targeted brain region—the left pACC, right pACC, left cOFC, and right cOFC—a series of six images is presented. Each series begins and ends with an image of an intact brain section, providing anatomical context, with four intermediate images demonstrating the electrode tracks within the tissue. Electrode tracks are clearly visible as linear disruptions in the cortical tissue. The inclusion of intact brain sections at the beginning and end of each series provides anatomical landmarks and orientation, facilitating precise localization of electrode placements within the brain.

**Figure S3.**
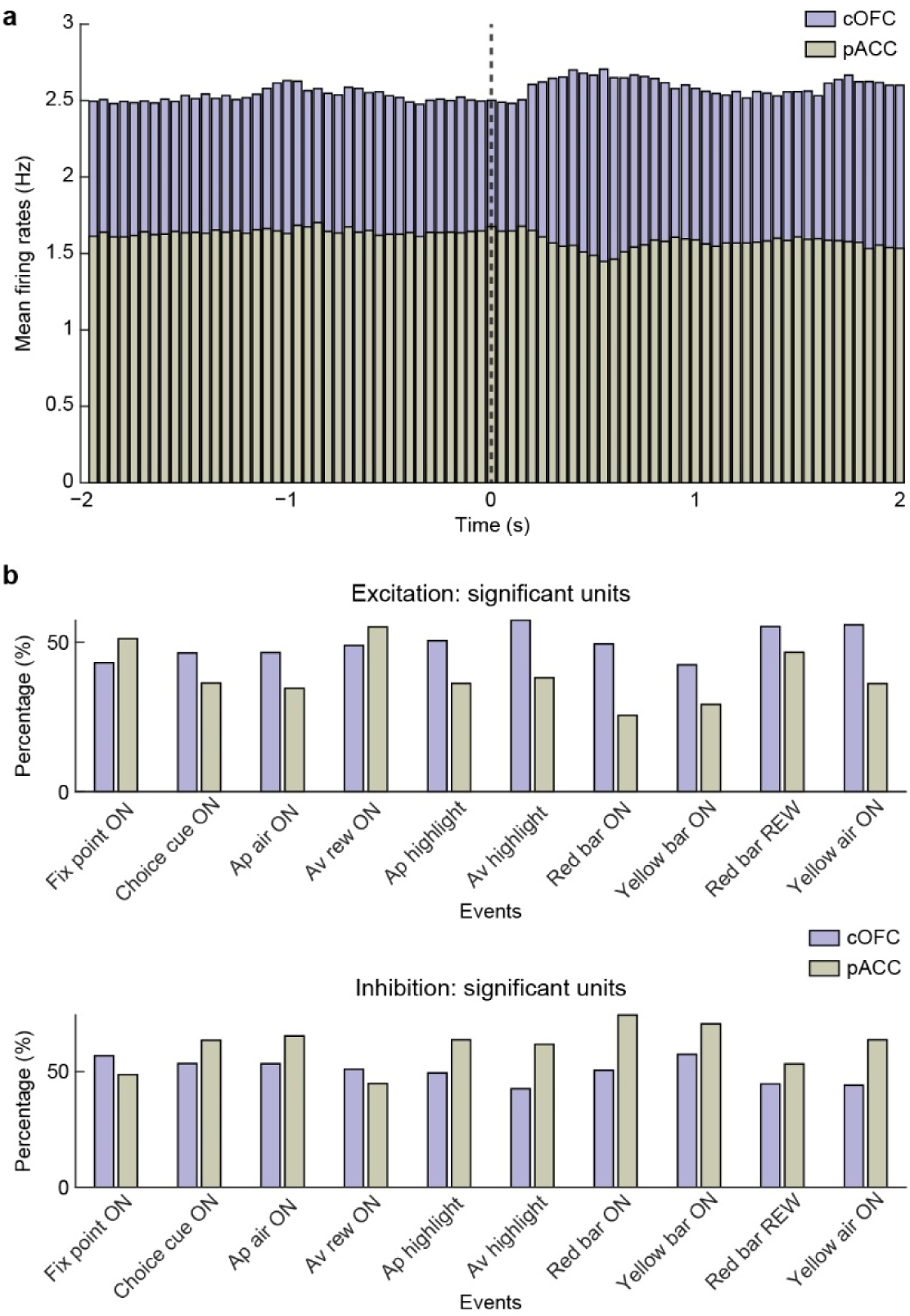
Excitation-Inhibition during Ap-Av task. **a,** Mean firing rates of all units with significant responses in the cOFC and pACC around the choice cue onset event (2 s before and after) of the Ap-Av task, with the black line in the middle indicating the onset of the event. Higher firing rates were observed for the cOFC compared to the pACC. Additionally, there was noticeable excitation in the cOFC after the onset of the cue and slight inhibition in the pACC. **b,** The proportion of units with statistically significant excitatory (top) and inhibitory (bottom) responses, calculated in relation to each event (Fix point ON: fixation point onset, Choice cue ON: choice cue onset, Ap air ON: approach airpuff onset, Av rew ON: avoidance reward onset, Ap highlight: approach highlight onset around cross, Av highlight: avoidance highlight onset around square, Red bar ON: Pavlovian red bar onset, Yellow bar ON: Pavlovian yellow bar onset, Red bar REW: Pavlovian red bar reward delivery, Yellow air ON: Pavlovian yellow bar airpuff delivery). The cOFC had a larger percentage of excitation (80% of the events) compared to the pACC (20% of the events). The exact opposite pattern was observed for the significant inhibition units.

**Figure S4.**
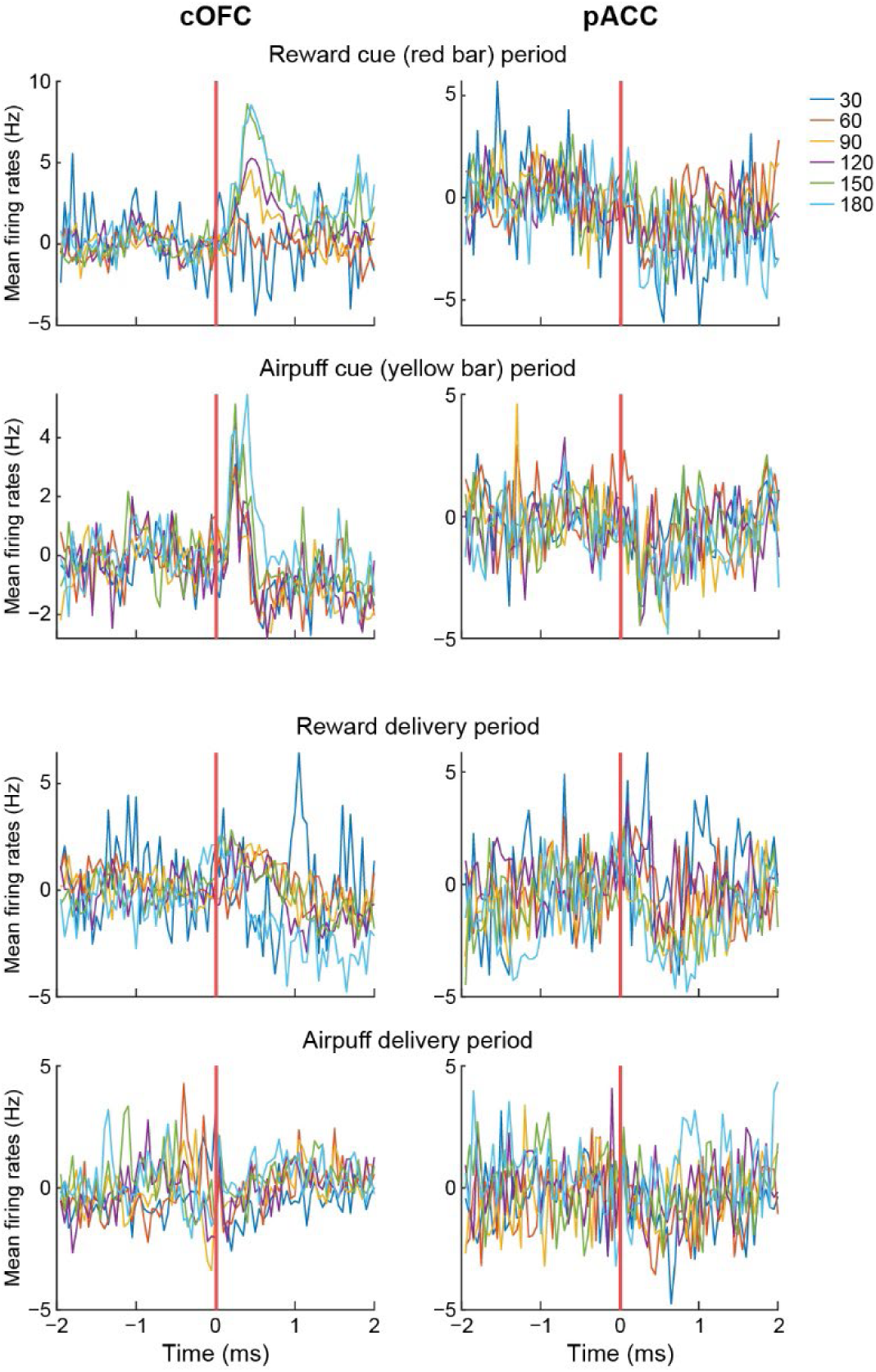
Mean firing rate of cOFC and pACC units recorded during Pavlovian trials. Each event’s onset is marked by a red vertical line at the midpoint of the x-axis in the plots, encompassing 2 s before and 2 s after the event onset. The left column represents data from the cOFC, broken down into the Cue and Outcome periods (2 panels each for both types of stimuli/outcomes). The right column shows the corresponding results from pACC. The different line colors indicate the different sizes of the red or yellow bar and the different durations of the actual outcome (reward or airpuff). Data presented include all recorded units and sessions.

**Figure S5.**
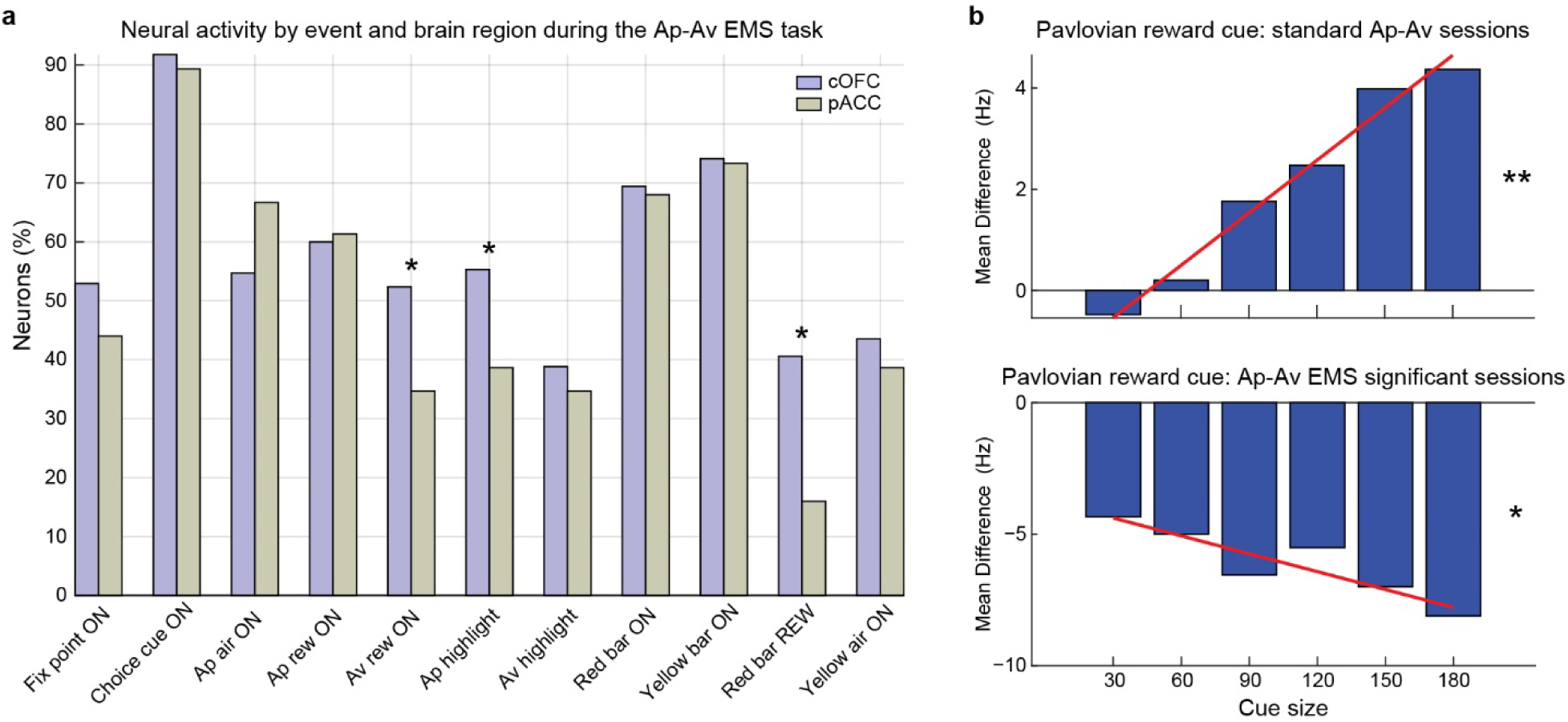
Event-related neural activity during Ap-Av EMS. **a,** Recordings were made from 245 well-isolated single units in the cOFC (170 units) and the pACC (75 units) of monkeys P and D. The percentage of neurons that exhibited statistically significant effects (chi-square test, p < 0.05, testing the change in spike counts 1 s before and 1 s after the onset of each event) during various task events across the two regions of interest was plotted. Percentage of statistically significant units was increased across all cue-related events (i.e., Choice cue ON, Red bar ON, Yellow bar ON) when EMS was applied (both stimulation levels), in comparison with the standard Ap-Av task without EMS. Interestingly, the cOFC appeared to be marginally more active across most task events, similar to the Ap-Av task. However, in two instances, the opposite pattern was observed, specifically during the approach trials outcome delivery events: approach airpuff onset (Ap air ON) and approach reward onset (Ap rew ON). Among the events analyzed, these three events—avoidance reward onset (Av rew ON; chi-square = 6.54, p = 0.0106), red bar reward onset (Red bar REW; chi-square = 14.22, p = 0.0002) and approach highlight onset (Ap highlight; chi-square = 5.76, p = 0.0164)—exhibited a statistically significant difference in neuronal activity between the cOFC and pACC. Thus, during EMS, a higher proportion of cOFC neurons showed significant activity than pACC neurons, indicating a potentially greater involvement or sensitivity of the cOFC towards positive outcomes during reward-related events. Also, the statistically significant effects observed during the cost/airpuff, which previously favored the pACC over the cOFC, were no longer present. Data were aggregated from both high and low current EMS sessions. **b,** Significant parametric modulation of Pavlovian Reward Cue firing rates in the cOFC during standard Ap-Av sessions (t(4) = 14.21, **p < 0.0001; top panel) and during Ap-Av EMS sessions with cOFC EMS (t(4) = −4.48, *p = 0.011; bottom panel).

**Figure S6.**
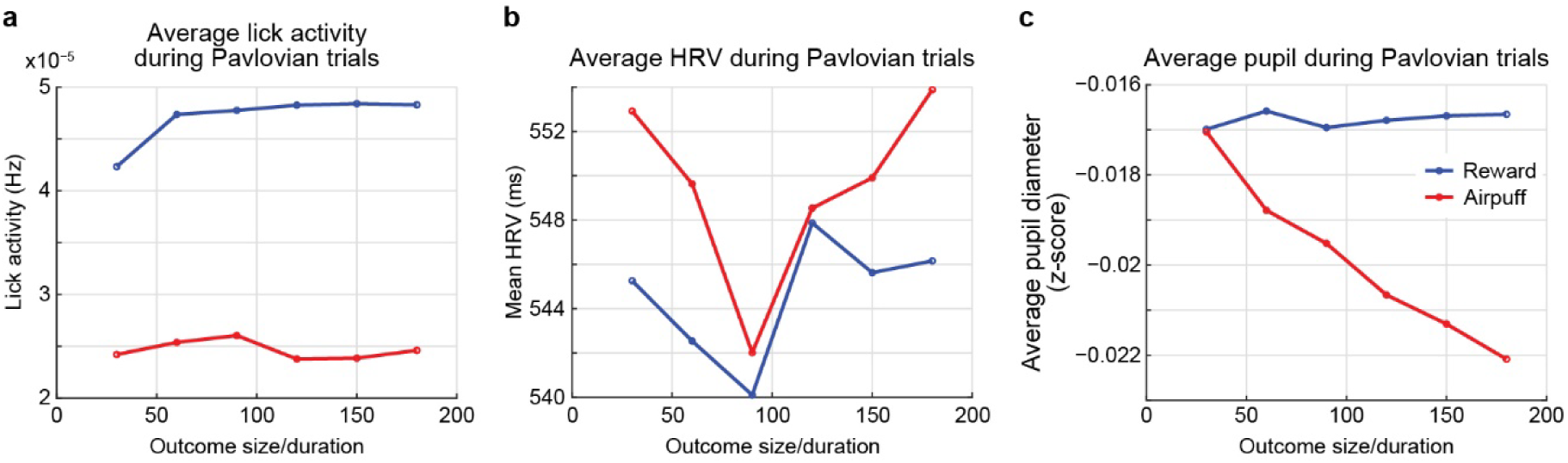
Physiological metrics during Pavlovian trials in the Ap-Av task. All metrics were averaged over the time window starting from cue presentation and ending just before the outcome period. **a,** Lick activity during Pavlovian trials, plotted against varying outcome size/duration for reward (blue) and airpuff (red) trials. **b,** Mean HRV during Pavlovian trials for reward and airpuff conditions, showing differential responses to increasing outcome sizes. **c,** Average pupil diameter (z-score) for reward and airpuff trials as a function of outcome size/duration. These metrics highlight distinct physiological responses depending on the type and size of the anticipated outcome in Pavlovian trials.

**Figure S7.**
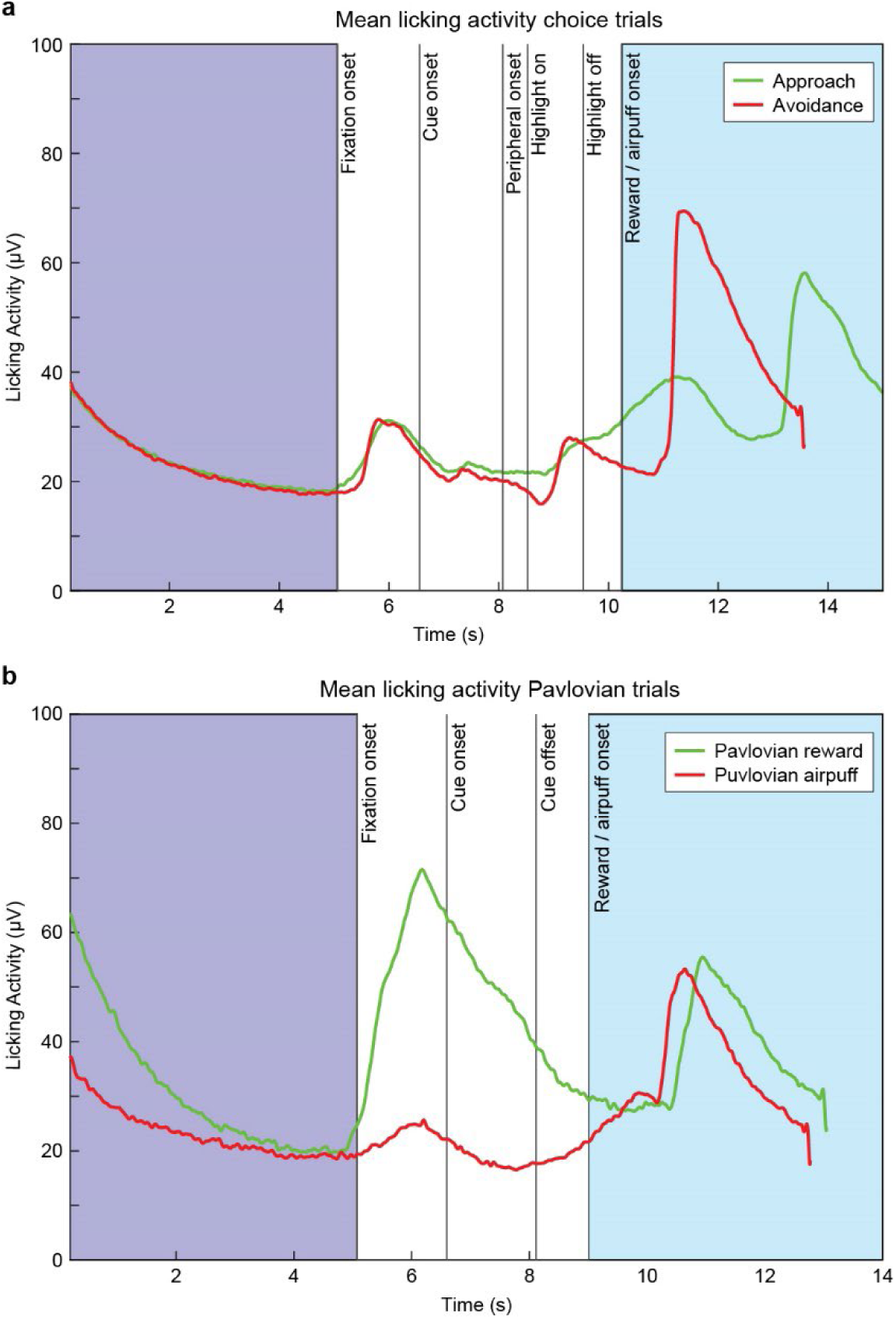
Licking activity during the Ap-Av task. The grand mean licking activity as a function of trial time across all trials in all Ap-Av sessions plotted for both the choice (**a**) and Pavlovian (**b**) trials. In the choice trials (**a**), it was observed that from the fixation onset, there was an equal increase in licking activity for both trial types, as the monkeys had not yet received information about the offer. As the offer was presented, slightly elevated activity was noted for the trials that would end in approach (green color), which continued to increase until the outcome periods. During the outcome periods, elevated licking activity was observed at different time points that coincided with the delivery of the rewards. For the Pavlovian trials (**b**), a remarkable elevation in licking activity was observed during the fixation point for trials where the monkey would freely receive a reward (green color).

**Figure S8.**
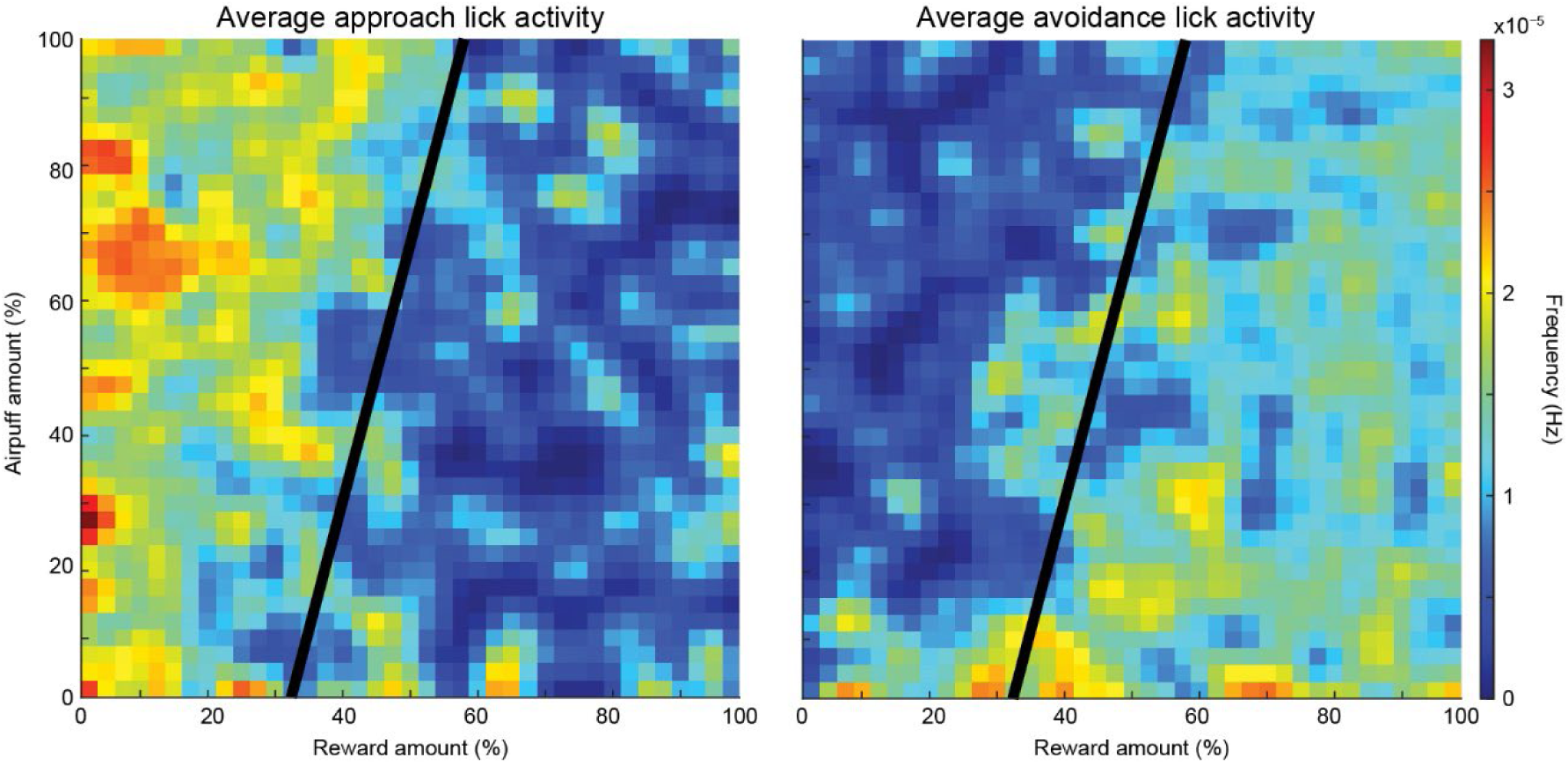
Lick activity during Ap and Av trials across varying reward and airpuff amounts. Heatmaps display the average lick activity (Hz) for Ap (left) and Av (right) trials. Lick activity is color-coded, with cooler colors (blue) indicating lower lick activity and warmer colors (yellow/red) indicating higher lick activity. During Ap trials, lick activity was high in conditions with high airpuff and small rewards, suggesting increased anticipation or anxiety when outcomes involve high costs. In Av trials, lick activity was lower in conditions with high airpuff and small rewards, reflecting a decisive rejection of unfavorable outcomes, whereas higher lick activity appears when airpuff levels are low and reward levels are high, indicating reduced aversion and increased anticipation in these conditions.

## References

Aggleton, J. P., Wright, N. F., Rosene, D. L., & Saunders, R. C. (2015). Complementary Patterns of Direct Amygdala and Hippocampal Projections to the Macaque Prefrontal Cortex. Cerebral cortex (New York, N.Y. : 1991), 25(11), 4351–4373. 10.1093/cercor/bhv019

Appelhans, B. M., & Luecken, L. J. (2006). Heart rate variability as an index of regulated emotional responding. Review of General Psychology, 10(3), 229–240. 10.1037/1089-2680.10.3.229

Amemori, K., & Graybiel, A. M. (2012). Localized microstimulation of primate pregenual cingulate cortex induces negative decision-making. Nature neuroscience, 15(5), 776–785. 10.1038/nn.3088

Amemori, K. I., Amemori, S., Gibson, D. J., & Graybiel, A. M. (2018). Striatal Microstimulation Induces Persistent and Repetitive Negative Decision-Making Predicted by Striatal Beta-Band Oscillation. Neuron, 99(4), 829–841.e6. 10.1016/j.neuron.2018.07.022

Amemori, S., Amemori, K. I., Yoshida, T., Papageorgiou, G. K., Xu, R., Shimazu, H., Desimone, R., & Graybiel, A. M. (2020). Microstimulation of primate neocortex targeting striosomes induces negative decision-making. The European journal of neuroscience, 51(3), 731–741. 10.1111/ejn.14555

Amemori S, Graybiel AM and Amemori K-i (2021) Causal Evidence for Induction of Pessimistic Decision-Making in Primates by the Network of Frontal Cortex and Striosomes. Front. Neurosci. 15:649167. doi: 10.3389/fnins.2021.649167

Amemori, S., Graybiel, A. M., & Amemori, K. I. (2024). Cingulate microstimulation induces negative decision-making via reduced top-down influence on primate fronto-cingulo-striatal network. Nature communications, 15(1), 4201. 10.1038/S41467-024-48375-1

Aron, A. R., Robbins, T. W., & Poldrack, R. A. (2014). Inhibition and the right inferior frontal cortex: one decade on. Trends in cognitive sciences, 18(4), 177–185. 10.1016/j.tics.2013.12.003

Aupperle RL, Melrose AJ, Francisco A, Paulus MP, and Stein MB (2015). Neural substrates of approach-avoidance conflict decision-making. Hum Brain Mapp 36, 449–462.

Ballesta, S., Shi, W., Conen, K. E., & Padoa-Schioppa, C. (2020). Values encoded in orbitofrontal cortex are causally related to economic choices. Nature, 588(7838), 450–453. 10.1038/S41586-020-2880-x

Barbas, H., & Rempel-Clower, N. (1997). Cortical structure predicts the pattern of corticocortical connections. Cerebral cortex (New York, N.Y. : 1991), 7(7), 635–646. 10.1093/cercor/7.7.635

Bechara, A., Damasio, H., & Damasio, A. R. (2000). Emotion, decision making and the orbitofrontal cortex. Cerebral cortex (New York, N.Y. : 1991), 10(3), 295–307. 10.1093/cercor/10.3.295

Bradley, M. M., Miccoli, L., Escrig, M. A., & Lang, P. J. (2008). The pupil as a measure of emotional arousal and autonomic activation. Psychophysiology, 45(4), 602–607. 10.1111/j.1469-8986.2008.00654.x

Chau, B. K., Sallet, J., Papageorgiou, G. K., Noonan, M. P., Bell, A. H., Walton, M. E., & Rushworth, M. F. (2015). Contrasting Roles for Orbitofrontal Cortex and Amygdala in Credit Assignment and Learning in Macaques. Neuron, 87(5), 1106–1118. 10.1016/j.neuron.2015.08.018

Chrysikou, E. G., Wing, E. K., & van Dam, W. O. (2022). Transcranial Direct Current Stimulation Over the Prefrontal Cortex in Depression Modulates Cortical Excitability in Emotion Regulation Regions as Measured by Concurrent Functional Magnetic Resonance Imaging: An Exploratory Study. Biological psychiatry. Cognitive neuroscience and neuroimaging, 7(1), 85–94. 10.1016/j.bpsc.2019.12.004

Critchley, H. D., Elliott, R., Mathias, C. J., & Dolan, R. J. (2000). Neural activity relating to generation and representation of galvanic skin conductance responses: a functional magnetic resonance imaging study. The Journal of neuroscience : the official journal of the Society for Neuroscience, 20(8), 3033–3040. 10.1523/JNEUROSCI.20-08-03033.2000

Damasio A. R. (1996). The somatic marker hypothesis and the possible functions of the prefrontal cortex. Philosophical transactions of the Royal Society of London. Series B, Biological sciences, 351(1346), 1413–1420. 10.1098/rstb.1996.0125

Drevets, W. C., Price, J. L., & Furey, M. L. (2008). Brain structural and functional abnormalities in mood disorders: implications for neurocircuitry models of depression. Brain structure & function, 213(1-2), 93–118. 10.1007/s00429-008-0189-x

Eblen, F., & Graybiel, A. M. (1995). Highly restricted origin of prefrontal cortical inputs to striosomes in the macaque monkey. The Journal of neuroscience : the official journal of the Society for Neuroscience, 15(9), 5999–6013. 10.1523/JNEUROSCI.15-09-05999.1995

Etkin, A., Egner, T., & Kalisch, R. (2011). Emotional processing in anterior cingulate and medial prefrontal cortex. Trends in cognitive sciences, 15(2), 85–93. 10.1016/j.tics.2010.11.004

Folkedal, O., Stien, L. H., Torgersen, T., Oppedal, F., Olsen, R. E., Fosseidengen, J. E., Braithwaite, V. A., & Kristiansen, T. S. (2012). Food anticipatory behaviour as an indicator of stress response and recovery in Atlantic salmon post-smolt after exposure to acute temperature fluctuation. Physiology & behavior, 105(2), 350–356. 10.1016/j.physbeh.2011.08.008

Feingold, J., Desrochers, T. M., Fujii, N., Harlan, R., Tierney, P. L., Shimazu, H., Amemori, K., & Graybiel, A. M. (2012). A system for recording neural activity chronically and simultaneously from multiple cortical and subcortical regions in nonhuman primates. Journal of neurophysiology, 107(7), 1979–1995. 10.1152/jn.00625.2011

Fouragnan, E. F., Chau, B. K. H., Folloni, D., Kolling, N., Verhagen, L., Klein-Flügge, M., Tankelevitch, L., Papageorgiou, G. K., Aubry, J. F., Sallet, J., & Rushworth, M. F. S. (2019). The macaque anterior cingulate cortex translates counterfactual choice value into actual behavioral change. Nature neuroscience, 22(5), 797–808. 10.1038/S41593-019-0375-6

Friedman, A., Homma, D., Bloem, B., Gibb, L. G., Amemori, K. I., Hu, D., Delcasso, S., Truong, T. F., Yang, J., Hood, A. S., Mikofalvy, K. A., Beck, D. W., Nguyen, N., Nelson, E. D., Toro Arana, S. E., Vorder Bruegge, R. H., Goosens, K. A., & Graybiel, A. M. (2017). Chronic Stress Alters Striosome-Circuit Dynamics, Leading to Aberrant Decision-Making. Cell, 171(5), 1191–1205.e28. 10.1016/j.cell.2017.10.017

Ghashghaei, H. T., Hilgetag, C. C., and Barbas, H. (2007). Sequence of information processing for emotions based on the anatomic dialogue between prefrontal cortex and amygdala. Neuroimage 34, 905–923. doi: 10.1016/j.neuroimage.2006.09.046

Godlewska, B. R., Norbury, R., Selvaraj, S., Cowen, P. J., & Harmer, C. J. (2012). Short-term SSRI treatment normalises amygdala hyperactivity in depressed patients. Psychological medicine, 42(12), 2609–2617. 10.1017/S0033291712000591

Gygax, L., Reefmann, N., Wolf, M., & Langbein, J. (2013). Prefrontal cortex activity, sympatho-vagal reaction and behaviour distinguish between situations of feed reward and frustration in dwarf goats. Behavioural brain research, 239, 104–114. 10.1016/j.bbr.2012.10.052

Ironside, M., Amemori, K. I., McGrath, C. L., Pedersen, M. L., Kang, M. S., Amemori, S., Frank, M. J., Graybiel, A. M., & Pizzagalli, D. A. (2020). Approach-Avoidance Conflict in Major Depressive Disorder: Congruent Neural Findings in Humans and Nonhuman Primates. Biological psychiatry, 87(5), 399–408. 10.1016/j.biopsych.2019.08.022

Hwang J, Mitz AR, Murray EA (2019) NIMH MonkeyLogic: Behavioral control and data acquisition in MATLAB. J Neurosci Methods 323:13–21. 10.1016/j.jneumeth.2019.05.002.

Kalin, N. H., Shelton, S. E., & Davidson, R. J. (1991). Defensive behaviors in infant rhesus monkeys: Ontogeny and context-dependent selective expression. Journal of Neuroscience, 11(11), 3064–3076.

Kalin, N. H., Shelton, S. E., & Davidson, R. J. (1996). Individual differences in freezing and cortisol in infant and mother rhesus monkeys. Behavioral Neuroscience, 110(3), 620–626.

Kringelbach, M. L., & Rolls, E. T. (2004). The functional neuroanatomy of the human orbitofrontal cortex: evidence from neuroimaging and neuropsychology. Progress in neurobiology, 72(5), 341–372. 10.1016/j.pneurobio.2004.03.006

Levy, D. J., & Glimcher, P. W. (2010). Neural systems for decision making. Wiley Encyclopedia of Operations Research and Management Science.

Levy I, Snell J, Nelson AJ, Rustichini A, Glimcher PW. Neural representation of subjective value under risk and ambiguity. J. Neurophysiol. 2010;103:1036–1047.

Mayberg H. S. (1997). Limbic-cortical dysregulation: a proposed model of depression. The Journal of neuropsychiatry and clinical neurosciences, 9(3), 471–481. 10.1176/jnp.9.3.471

McFadden D. Conditional logit analysis of qualitative choice behavior. In: Zarembka P, editor. Frontiers in econometrics. New York: Academic Press; 1974. pp. 105–142.

Millan MJ (2003). The neurobiology and control of anxious states. Prog Neurobiol 70, 83–244.

Mnih, V., Badia, A. P., Mirza, M., Graves, A., Lillicrap, T., Harley, T., Lillicrap, T.P., Silver, D., Kavukcuoglu, K. (2016). Asynchronous methods for deep reinforcement learning. International conference on machine learning, 1928–1937.

Murphy, P. R., Vandekerckhove, J., & Nieuwenhuis, S. (2014). Pupil-linked arousal determines variability in perceptual decision making. PLoS computational biology, 10(9), e1003854. 10.1371/journal.pcbi.1003854

Murray, E. A., & Fellows, L. K. (2022). Prefrontal cortex interactions with the amygdala in primates. Neuropsychopharmacology : official publication of the American College of Neuropsychopharmacology, 47(1), 163–179. 10.1038/S41386-021-01128-w

Nassar, M. R., Rumsey, K. M., Wilson, R. C., Parikh, K., Heasly, B., & Gold, J. I. (2012). Rational regulation of learning dynamics by pupil-linked arousal systems. Nature neuroscience, 15(7), 1040–1046. 10.1038/nn.3130

O’Doherty, J., Kringelbach, M. L., Rolls, E. T., Hornak, J., & Andrews, C. (2001). Abstract reward and punishment representations in the human orbitofrontal cortex. Nature neuroscience, 4(1), 95–102. 10.1038/82959

Padoa-Schioppa C. Neurobiology of economic choice: a good-based model. Annu. Rev. Neurosci. 2011;34:333–359.

Papageorgiou, G. K., Sallet, J., Wittmann, M. K., Chau, B. K. H., Schüffelgen, U., Buckley, M. J., & Rushworth, M. F. S. (2017). Inverted activity patterns in ventromedial prefrontal cortex during value-guided decision-making in a less-is-more task. Nature communications, 8(1), 1886. 10.1038/S41467-017-01833-5

Price, J. L., & Drevets, W. C. (2010). Neurocircuitry of mood disorders. Neuropsychopharmacology : official publication of the American College of Neuropsychopharmacology, 35(1), 192–216. 10.1038/npp.2009.104

Rempel-Clower, N. L., & Barbas, H. (2000). The laminar pattern of connections between prefrontal and anterior temporal cortices in the Rhesus monkey is related to cortical structure and function. Cerebral cortex (New York, N.Y. : 1991), 10(9), 851–865. 10.1093/cercor/10.9.851

Ressler, K. J., & Mayberg, H. S. (2007). Targeting abnormal neural circuits in mood and anxiety disorders: from the laboratory to the clinic. Nature neuroscience, 10(9), 1116–1124. 10.1038/nn1944

Rolls E. T. (2004). The functions of the orbitofrontal cortex. Brain and cognition, 55(1), 11–29. 10.1016/S0278-2626(03)00277-X

Robbins T. W. (2007). Shifting and stopping: fronto-striatal substrates, neurochemical modulation and clinical implications. Philosophical transactions of the Royal Society of London. Series B, Biological sciences, 362(1481), 917–932. 10.1098/rstb.2007.2097

Rushworth, M. F., & Behrens, T. E. (2008). Choice, uncertainty and value in prefrontal and cingulate cortex. Nature neuroscience, 11(4), 389–397. 10.1038/nn2066

Rushworth, M. F., Noonan, M. P., Boorman, E. D., Walton, M. E., & Behrens, T. E. (2011). Frontal cortex and reward-guided learning and decision-making. Neuron, 70(6), 1054–1069. 10.1016/j.neuron.2011.05.014

Shackman, A. J., Salomons, T. V., Slagter, H. A., Fox, A. S., Winter, J. J., & Davidson, R. J. (2011). The integration of negative affect, pain and cognitive control in the cingulate cortex. Nature reviews. Neuroscience, 12(3), 154–167. 10.1038/nrn2994

Sammito, S., & Thielmann, B. (2020). The circadian rhythm of heart rate variability. Biological Rhythm Research, 51(2), 186–195.

Thayer, J. F., & Lane, R. D. (2000). A model of neurovisceral integration in emotion regulation and dysregulation. Journal of affective disorders, 61(3), 201–216. 10.1016/s0165-0327(00)00338-4

Volkow, N. D., Wang, G. J., Fowler, J. S., & Tomasi, D. (2012). Addiction circuitry in the human brain. Annual review of pharmacology and toxicology, 52, 321–336. 10.1146/annurev-pharmtox-010611-134625

Von Neumann J, Morgenstern O. Theory of games and economic behavior. Princeton: Princeton Univ. Press; 1947.

Wallis J. D. (2007). Orbitofrontal cortex and its contribution to decision-making. Annual review of neuroscience, 30, 31–56. 10.1146/annurev.neuro.30.051606.094334

Xu, X., Dai, J., Chen, Y., Liu, C., Xin, F., Zhou, X., Zhou, F., Stamatakis, E. A., Yao, S., Luo, L., Huang, Y., Wang, J., Zou, Z., Vatansever, D., Kendrick, K. M., Zhou, B., & Becker, B. (2021). Intrinsic connectivity of the prefrontal cortex and striato-limbic system respectively differentiate major depressive from generalized anxiety disorder. Neuropsychopharmacology : official publication of the American College of Neuropsychopharmacology, 46(4), 791–798. 10.1038/S41386-020-00868-5

